# Differentiation between wild and domesticated Ungulates based on ecological niches

**DOI:** 10.1101/629188

**Authors:** Elke Hendrix, Rutger Vos

## Abstract

The domestication of flora and fauna is one of the most significant transitions in humankind’s history. It changed human societies drastically with alterations in biodiversity, atmospheric composition and land use. Humans have domesticated relatively few large animals and all of them belong to the Ungulates, though they are only 15 species of the ±150 that the entire group comprises. This can partially be explained by behavioral and life history pre-adaptations, e.g. social group structure, mating behavior, parent-young interaction, feeding behavior, and response to humans. The other dimension of proposed pre-adapatations concerns the biomes from which domesticated Ungulates originate. Here we test whether environmental preferences i.e. niches and related niche traits, differentiate between wild and domesticated Ungulates. We used three methods to determine the niche dimensions for each species and calculate overlap in niche space between them. Two methods are based on MaxEnt ecological niche models and one method uses raw occurrence data. Our results show that there is no weighted combination of environmental traits that clusters all domesticated Ungulates to the exclusion of all wild ones. On the contrary, domesticated Ungulates are overdispersed in niche space, indicating that the major pre-adaptations for domestication are not directly related to the abiotic niche. However, phylogenetic generalized linear modelling of selected niche dimensions does predict domestication significantly. We conclude that further research of other traits is needed.

## Introduction

The domestication of flora and fauna is one of the most significant transitions in humankind’s history and domination of planet earth (1, 2). Domestication can be understood as the alteration of wild species by selecting traits that are useful to humans. Examples of this are the selection of dogs that are able to live with people or the selection of wheat that have more seeds per plant (1). The first known animal domestication is the domestication of the wolf between 17.000 and 15.000 years BP in the Old World (3, 4). Dogs were later followed by goats and sheep that were domesticated first in southern and southwestern Asia between 11.000 and 9.000 BP (5, 6). Pigs and cattle were also domesticated in southwestern Asia around 8.000 BP, followed by horses and camels around 6.000 BP in central Asia (4). Some of these species were later domesticated independently in other parts of the earth (7, 8).

Domesticated flora and fauna interacted and increased food production; for example, large mammals were used to pull plows and provide manure, making previously un-farmable areas suitable for domesticated plants. Large mammals were also used as a source of milk to produce butter and cheese, yielding much more calories in their lifetime than if they were only used for meat production (9). The food surpluses meant that not everyone had to be involved in the production of food eventually creating extra time for new developments and innovations. Domestication and the agricultural economies based on them changed human societies drastically with alterations in biodiversity, land use and atmospheric compositions (10).

Domestication of flora and fauna did not only alter human societies but it changed the domesticates to the extent that they greatly differ from their wild ancestors (2). In general, domesticates have a set of common traits which are called the “domestication syndrome”. For plants, some of these common traits include that they are able to produce bigger seeds, the ability to reduce physical and chemical defences and the reduced ability to disperse their seeds without the intervention of humans (2, 11–13). For animals the domestication syndrome exhibits itself in floppy ears, an increase in docile behaviour, size reduction, and facial neotony (12).

Over the course of thousands of years, humans have domesticated relatively few large animals. The group of large hoofed mammals, which we refer to here by the folk-taxonomic “Ungulates” (i.e. terrestrial Perissodactyla + Artiodactyla), include species such as donkeys, cattle, pigs and giraffes. We have only domesticated about 15 Ungulates species from the +− 150 Ungulates. Interestingly, some of the 15 domesticated Ungulate species were domesticated multiple times independently through space and time, while the other +150 Ungulate species were never successfully domesticated, for a variety of reasons. Some of these reasons can be explained by behavioral pre-adaptations like social structure, sexual behaviour, parent-young interaction, feeding behaviour, and response to humans and new environments (14). (1) Animals that live in herds with a hierarchical social structures are the most suited for domestication because humans can effectively take over the top rank in the hierarchy. (2) Also sexual behaviour is of critical importance since some animal species do not breed in captivity, e.g. cheetahs. (3) The speed at which parents bond with their offspring needs to be fast and created through imprinting. (4) Feeding behaviour determines whether it is efficient to domesticate a specific species. (5) The response to humans is of importance because some animals have a tremendous flight response while others, e.g. zebras, have unpredictable and aggressive temperaments (9).

An untested theory is that next to behavioral pre-adaptations, the biomes where the Ungulates originate from determine the ability of a species to be domesticated. According to ecological theory, each species has a niche within an ecosystem where the environmental preferences i.e. niche traits, are according to individual fitness (15–18). Hutchinson (1957) described a niche as a n-dimensional hypervolume where the dimensions are explained by environmental variables e.g. temperature and rainfall (16). Here we use the term niche traits to describe the unique set of mean environmental variables per species.

Niche space and related niche traits can be captured by numerical models called Ecological Niche Models (ENM), which link species distributions to their environment (19). The use of ENMs has been applied in a wide variety of fields, such as evolutionary biology and conservation biology, for example to map the probable distribution of species after climate change (20). ENMS are empirical models that correlate environmental data, e.g. climate data and soil characteristics, to occurrence locations to identify the environmental conditions needed to maintain populations (20, 21). The result of an ENM can be, for example, that species × has only been recorded in environmental conditions in which the soil pH is lower than 7 and the precipitation is between 60 mm and 120mm. The suitable environmental conditions can be used to map suitable sites for species × all over the world. Hence, it should be noted that suitable areas (fundamental niche) and the actual occurrence (realized niche) of species is not the same: geographic barriers or biotic interactions can limit the dispersal to suitable sites (20).

The comparison of Hutchinsonian niche space per species can give some interesting insight in overlap or differences in niche space and related niche traits. For this research we use ENMs and raw occurrence data to analyse whether there is a significant difference in habitats for domesticated ungulates and wild ungulates.

## Materials

All the source code, workflows, and data sets used in this research can be found and downloaded at our GitHub repository (https://github.com/naturalis/trait-geo-diverse-ungulates).

### A. Species occurrence data

Ungulates are a diverse group of hoofed placental mammals distributed over Afro-Eurasia and the New World. For this research, we extracted all available Ungulates occurrence data from the global biodiversity information facility (GBIF), which is a portal that collates both observation and digitized museum collection data (22, 23). Records in the GBIF database (exported as DarwinCore archives (24)) contain information about the type of occurrence, the taxon, the location in longitude and latitude, and various metadata. We selected a total of 152 Ungulates species, of which 14 domesticated species and 138 wild species. We used wild ancestors of domesticated species as a proxy because most of the domesticates live in captivity and are changed to such an extent that their preferences hardly resemble those of their wild ancestors. Therefore, any inferred niche dimensions of current, domesticated Ungulate species do not correctly portray their original niche preferences (see table 1 for a list of the wild ancestors). We thus take the wild ancestors (or proxies thereof) to model the original niches in which domestication took place.

**Table 1.**
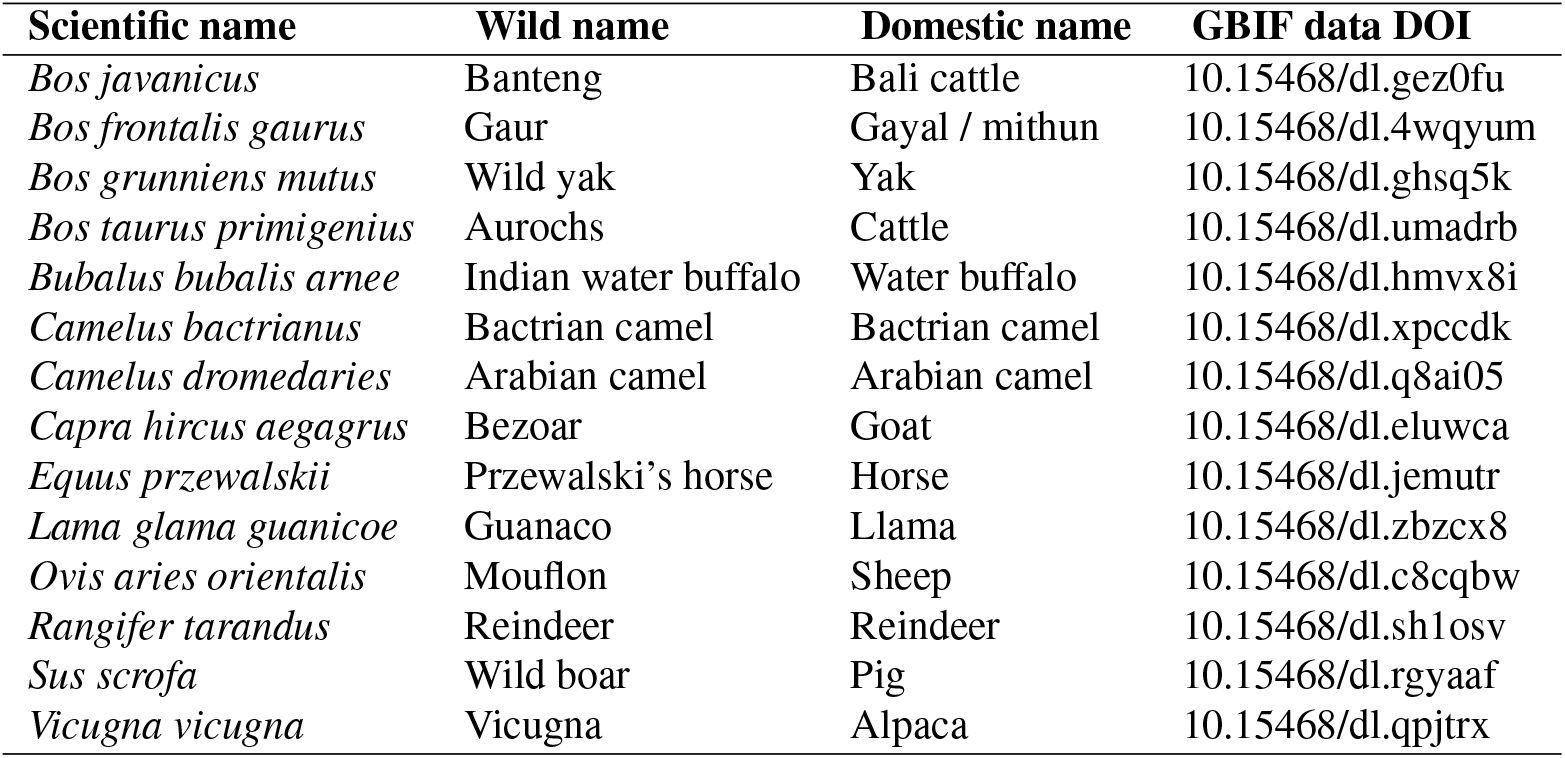
Wild ancestors of domesticated Ungulates

We collected occurrence data sets from GBIF using higher taxon searches for terrestrial Artiodactyla (even-toed ungulates), and Perissodactyla (odd-toed ungulates). The results of these searches are shown in table 2. To resolve taxonomic ambiguities in the records and to anchor the data on canonical names we downloaded the GBIF backbone taxonomy (25). From this we wrote all synonyms and non-accepted taxonomic names to a taxonomic name variants table that links to a canonical names table based on the backbone. We loaded these tables into a SQLite database to form our taxonomic name management structure, which we further reconciled with the *Mammal Species of the World* taxonomy (26) and for which we collected additional name variants from ITIS (27). We extracted the following columns from the DarwinCore occurrence archives: *gbifID, type, basisOfRecord, eventDate, decimalLatitude, decimalLongitude, datasetKey, hasGeospatialIssues and Taxonkey*. We loaded these columns into SQLite and anchored the records to the taxonomic backbone with the Taxonkey. We removed all the occurrence records with incomplete longitude, latitude and event-date fields, records with geospatial issues (*sensu* GBIF) and records of which the basis (human observation, fossil or preserved specimen) was unknown. To reduce the amount of spatial outliers in the data sets we only used species records that fall within the IUCN species distribution polygons (28) and whose mean pairwise distance to all others does not differ from the species mean by more than 1 standard deviation. To be able to compare the taxon names of species in the IUCN data set to the names in our data sets the names in the IUCN data set were reconciled to the aforementioned GBIF backbone taxonomy (25). Afterwards the records per separate species were exported to CSV files; only spatially distinct occurrences were retained, and only species with more than 10 records (29) were exported.

**Table 2.** GBIF taxon searches

| Taxon | Date | Occurrences | Datasets | DOI |
| --- | --- | --- | --- | --- |
| Artiodactyla | 2018-10-19 | 1,221,109 | 1,019 | https://doi.org/10.15468/dl.qqwyhp |
| Perissodactyla | 2018-10-20 | 226,805 | 289 | https://doi.org/10.15468/dl.jxwvia |

The data obtaining and cleaning workflow failed to produce sufficient records for a few of the wild species. Using custom queries, more data were obtained for *Budorcas taxicolor*, *Madoqua saltiana*, *Neotragus batesi*, *Ovis ammon*, *Procapra picticaudata*, *Rucervus duvaucelii*, *Tragulus javanicus* and *Tragulus kanchil*. We filtered the CSV files with species that were selected using custom queries in the R software environment(30): erroneous coordinates were removed using the *CoordinateCleaner* package, by removing sea coordinates, records near GBIF/biodiversity institutes and outliers (30, 31).

### B. GIS data

The environmental data GIS layers used in this research relate to climate, topography and soil characteristics. Climatic information was extracted from the widely used Bioclim data set, which includes 19 bioclimatic variables. The GIS layers contain information such as precipitation in the driest quarter, or maximum temperatures of the coldest month, and are constructed based on monthly remote sensing data between 1950 and 2000, with a spatial resolution of 5 minutes (table 3) (32).

**Table 3.**
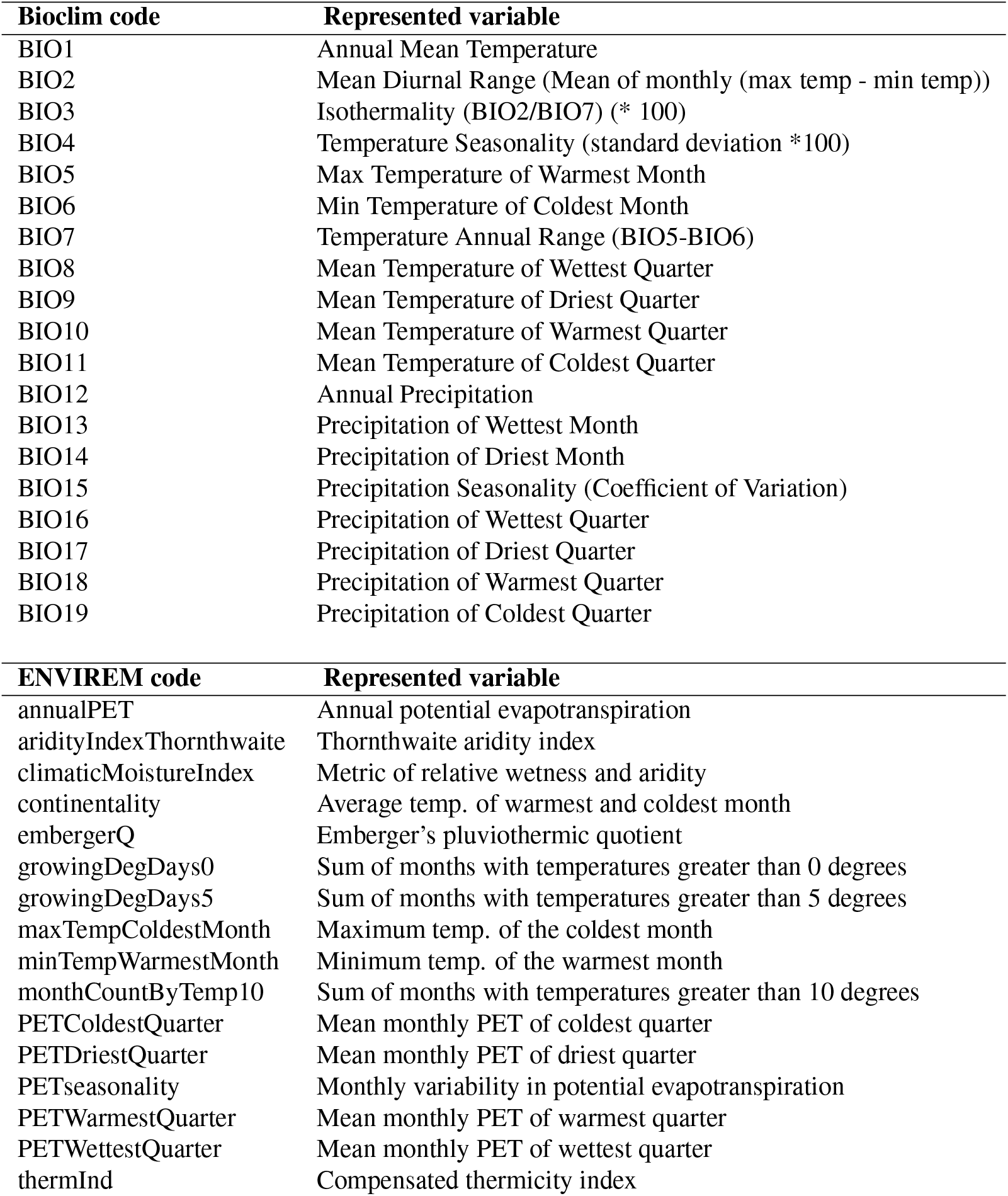
Bioclim (32) and ENVIREM datasets (33).

| Bioclim code | Represented variable |
| --- | --- |
| BIO1 | Annual Mean Temperature |
| BIO2 | Mean Diurnal Range (Mean of monthly (max temp - min temp)) |
| BIO3 | Isothermality (BIO2/BIO7) (* 100) |
| BIO4 | Temperature Seasonality (standard deviation *100) |
| BIO5 | Max Temperature of Warmest Month |
| BIO6 | Min Temperature of Coldest Month |
| BIO7 | Temperature Annual Range (BIO5-BIO6) |
| BIO8 | Mean Temperature of Wettest Quarter |
| BIO9 | Mean Temperature of Driest Quarter |
| BIO10 | Mean Temperature of Warmest Quarter |
| BIO11 | Mean Temperature of Coldest Quarter |
| BIO12 | Annual Precipitation |
| BIO13 | Precipitation of Wettest Month |
| BIO14 | Precipitation of Driest Month |
| BIO15 | Precipitation Seasonality (Coefficient of Variation) |
| BIO16 | Precipitation of Wettest Quarter |
| BIO17 | Precipitation of Driest Quarter |
| BIO18 | Precipitation of Warmest Quarter |
| BIO19 | Precipitation of Coldest Quarter |

| ENVIREM code | Represented variable |
| --- | --- |
| annualPET | Annual potential evapotranspiration |
| aridityIndexThornthwaite | Thornthwaite aridity index |
| climaticMoistureIndex | Metric of relative wetness and aridity |
| continentality | Average temp. of warmest and coldest month |
| embergerQ | Emberger's pluviothermic quotient |
| growingDegDays0 | Sum of months with temperatures greater than 0 degrees |
| growingDegDays5 | Sum of months with temperatures greater than 5 degrees |
| maxTempColdestMonth | Maximum temp. of the coldest month |
| minTempWarmestMonth | Minimum temp. of the warmest month |
| monthCountByTemp10 | Sum of months with temperatures greater than 10 degrees |
| PETColdestQuarter | Mean monthly PET of coldest quarter |
| PETDriestQuarter | Mean monthly PET of driest quarter |
| PETseasonality | Monthly variability in potential evapotranspiration |
| PETWarmestQuarter | Mean monthly PET of warmest quarter |
| PETWettestQuarter | Mean monthly PET of wettest quarter |
| thermInd | Compensated thermicity index |

Bioclim GIS layers with a spatial resolution of 2.5 minutes are also available, but we use the coarser data set because the GBIF occurrence data typically have an average error margin of +− 10 km anyway. Besides the Bioclim data we used the ENVIronmental Rasters for Ecological Modelling (ENVIREM) GIS layers. These contain 16 additional climate variables complementary to the Bioclim data set and are directly relevant to the physiological and ecological processes that determine the distribution of species (table 3) (33).

Median elevation variables were extracted from the Harmonized World Soil Database (HWSD) and are based on NASA’s Shuttle Radar Topographic Mission (SRTM) dataset. The HWSD median elevation data set provides layers with a 30 second spatial resolution. The topographic wetness index and the terrain roughness index are extracted from the ENVIREM dataset and have a spatial resolution of 30 seconds (33). The median elevation variables were used to calculate both slope and aspect also in the R software environment using the *terrain* function in the *raster* package (34). Slope and aspect were calculated because elevation variables are directly correlated to temperature variables, and therefore should not be used directly (29).

We extracted bulk density, clay percentage, pHCaCL and organic carbon of the top layer (0 till 45 cm) from the Land-Atmosphere Interaction Research Group with a spatial resolution of 5 minutes (35). The bulk density, clay percentage, pHCaCL and organic carbon data sets were used because they are known to influence plant distribution and growth (36, 37) and are not highly correlated to one another (38).

The aforementioned environmental raster layers were resampled to the coarsest resolution (5 minutes) using the nearest neighbour sampling method in the R software environment (30).

## Methods

We used environmental data and species occurrences to calculate niche overlap using the following three methods.

- Niche trait overlap based on modelled habitat projections. Here we used the MaxEnt algorithm to derive the means of the environmental values within the projected suitable area of each species. We then used the niche traits to calculate the distance between the different species.
- Spatial niche overlap based on modelled habitat projections. The MaxEnt model projections were used to calculate overlap in niche space.
- Niche trait overlap based on raw occurrence points. We calculated the mean niche traits per species by directly extracting environmental variables on the occurrence locations. The mean niche traits per species were used to calculate the distance between the different species.

Two methods are based on overlap in niche traits and one methods is based on spatial overlap of habitats. The difference between the methods is that overlap based on niche traits is derived by comparing the mean environmental variables per species while the calculation of spatial overlap of habitats is based on the actual overlap of the niches projected on earth.

### C. Ecological niche modelling

To analyze the spatial distribution and related niche traits we used the Maximum Entropy (MaxEnt) machine learning algorithm version 3.3.3 to construct ENMs for the 152 filtered species. Previous research has demonstrated that the MaxEnt technique performs well when using occurrence only data to estimate the relationships between environmental predictors and the occurrence of species (39–42). The widely used machine learning algorithm is very efficient in the complex handling of response and predictor variable interactions and works well with little occurrence data points (39, 43, 44).

Before the construction of the MaxEnt model the environmental rasters are cropped to the extent of the occurrence points for a specific species + a buffer around the total extent. Buffers that are too small can result in underestimations of edge effects while buffer that are too large have the risk of losing track of favorable environmental conditions due to noise. In this research we used a buffer of 1.000 km around the occurrence points.

In order to avoid collinearity only uncorrelated cropped environmental raster layers can be used in the ENMs (29, 45). To remove correlated layers the *removeCollinearity* function in the *virtualspecies* version 1.4-4 R package was used (46). Environmental variables with a Pearson’s R correlation coefficients above 0.7 were grouped and one variable within this group was randomly chosen, resulting in a cropped raster stack with only uncorrelated environmental rasters. The cropped uncorrelated rasters and all available and filtered occurrence points are used in the ENM. The ENM is based on the MaxEnt algorithm and is constructed using the *MaxEnt* function from the *dismo* R package (47). The function extracts abiotic environmental data for the training occurrence locations and 1000 random sampled background locations, resulting in a model MaxEnt object that can be used to project worldwide which other locations are suitable for the species outside of the cropped model environment.

A common measure to assess the fit of MaxEnt models is the area under the receiver operating curve (ROC), also known as the AUC (48). The AUC can be interpreted as the probability that a randomly chosen “occurrence point”, the location of a species occurrence, has a higher predicted probability of occurrence than a randomly chosen absence point (29). Only ENMs with an AUC value higher than 0.7 are generally accepted as valid models (49). A mayor drawback of the use of AUC values to validate the models performance is that our models depend on background samples and not true absence samples. Therefore the maximum AUC value is not 1 but 1-*a*/2, where *a* is the true species distribution (29). The true distribution of the species is unknown, which makes the inter-comparison of species distribution models based on AUC values and the validation of AUC values impossible. Raes and ter Steege (2007) developed a null-model to overcome these challenges and test whether AUC values significantly deviate from random models (50). Here we constructed 100 random MaxEnt models based on randomly sampled occurrence points. If the true MaxEnt model performed better than 95% of the null-models the model was deemed valid.

To assess the effects of the different environmental data sets in the MaxEnt model we used the response curves and variable importance scores rendered by the *MaxEnt* function in the *dismo* package (47). Afterwards the valid models were used to project suitable niches per species outside of the clipped model environment. Projection values range from 0 (not suitable) to 1 (very suitable). We put thresholds on these values by removing the suitability values below the 10% worst performing occurrence points. The projections were saved with and without restrictions. The restrictions were based on Holt’s zoogeographical region, which divide the earth in 11 large realms (51). The zoogeographical realms with occurrence points are the limit of projections, because the areas outside of the realms are unlikely to be inhabitated by the species, due to biogeographical barriers to dispersal.

### D. Calculating overlap in niche trait space derived from modelled habitat projections

To assess whether ecological niche traits, based on MaxEnt niche projections restricted by the 11 zoogeographical realms, differ between domesticated and wild Ungulates, we calculate their niche trait values and overlap. The suitable areas in the restricted projection rasters were used to calculate the mean normalized environmental traits per species following a similar approach to the Outlying Mean Index (OMI) of Doledec et al. (2000) (52, 53). First we extracted all environmental variables in the suitable areas and standardized these values to a zero mean and a standard deviation of one by using the *dudi.pca* function in the *ade4* R package (54). Second we averaged the standardized values per species by using the *niche* function in the *ade4* R package (53, 54). The resulting dataframe with standardized environmental traits per species was used to calculate the overlap in niche traits using Gower’s distance metric. Gower’s distance is explained as follows by equation 1:

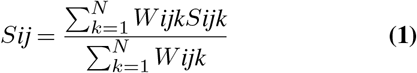

Where:

- S*ijk*: the distance between *i* and *j* for variable *k*
- W*ijk*: the weight that is given for variable *k* between two observations *i* and *j*

With Gower’s distance one can calculate the distance in environmental niche traits per species between all pairs of sample units (55). Species that have similar niche traits have a low Gower’s distance value and species that have different niche traits have a high Gower’s distance value. The mean environmental variable importance scores derived from the MaxEnt models were used to weigh the variable importance in the Gower’s distance calculations. We calculated Gower’s distance in the R software environment (30) with the *daisy* function in the cluster 2.0.7-1 R package (56).

### E. Calculating overlap in niche space based on modelled habitat projections

To assess whether ecological niches of domesticated ungulates and wild ungulates differ we calculate the spatial niche overlap by using the modelled habitat projections that are not restricted by the 11 realms. We used Schoener’s distance to calculate the distance in spatial niche overlap since it has been suggested to be the best suited index for MaxEnt ENM outputs (57, 58). We calculates Schoener’s index in the R software environment (30) with the *calc.niche.overlap* function in the *ENMeval* R package (59). The index ranges from 0 which is no overlap to 1 which is a complete overlap and is based on the model prediction maps. To create a measure over distance we subtracted the overlap, i.e. the inverse of the overlap from 1.

### F. Calculating overlap in niche trait space based on raw occurrence data

We calculated niche trait space overlap based on raw occurrence data by directly extracting niche traits from the environmental rasters underneath the occurrence points. Afterwards the niche traits were averaged and standardized per species following the OMI method proposed by Doledec et al. (2000) (52). The niche traits per species were used to calculate niche overlap with Gower’s distance metric and weights were given to the separate environmental variables.

### G. Clustering similar niches

The distance matrices based on (i) overlap in niche trait space derived from modelled habitat projections, (ii) overlap in niche space based on modelled habitat projections and (iii) overlap in niche trait space based on raw occurrence data were used to construct three dendrograms. The dendrograms were constructed in the R software environment (30) with the *as.phylo* function in the ape package (60). We calculated the optimal amount of clusters and grouped the similar species together. Afterwards we calculated the pairwise difference in distance between domesticated Ungulates and domesticated Ungulates and between domesticated Ungulates and wild Ungulates to see if domesticated Ungulates had a shorter distances and are therefore more similar.

### H. Predicting domestication based on niche traits

As a last step we constructed a generalized linear prediction model that predicts whether a species is domesticated or not based on the previously calculated niche traits (niche traits based on modelled habitat projections and niche traits based on raw occurrence data). For the construction of a generalized linear model independent observations are needed which is not the case for Ungulates. Most Ungulates are related and this needs to be corrected for in our model by adding the phylogenetic tree of present day mammals by Olaf et al. (2007) (61). We used a generalized linear model that corrects for dependent observations to construct a prediction model with the *phyloglm* function in the *phylolm* version 2.6 R package (62). The generalized linear prediction model was validated with Akaike’s information criterion (AIC) (63). AIC values give information about the relative quality of the prediction model. Actual AIC values are inconsequential and can only be used to compare values with (a model with the lowest AIC value is the model with the highest prediction accuracy) (64).

## Results

### I. Performance of ecological niche models

Figure 1 shows the restricted modelled habitat projections for the domesticated species. It is visible that for some species the projections remain close to the occurrence locations (red points) while for other locations the niche seems very wide and far away from the original occurrence locations. The *Bos grunniens mutus* (2) has occurrence points in Tibet, Mongolia and Russia but according to the modelled habitat projections parts of Canada and Greenland are also suitable. The ENM is mainly influenced by variables such as PET-WarmestQuarter, bio14 and aspect. The *Equus przewalskii* (9) also has occurrence points in Mongolia and shows the same projections in parts of Canada and Greenland. The most important variables in the ENM were bio1, bio14 and Bulk density. Lastly, the *Ovis aries orientalis* has occurrence points in Afghanistan, India and Kazakhstan and also shows projected suitable habitats in Canada. Variables that were important in the projection of suitable areas were bio14, PETwettestquarter and slope.

**Fig. 1.**
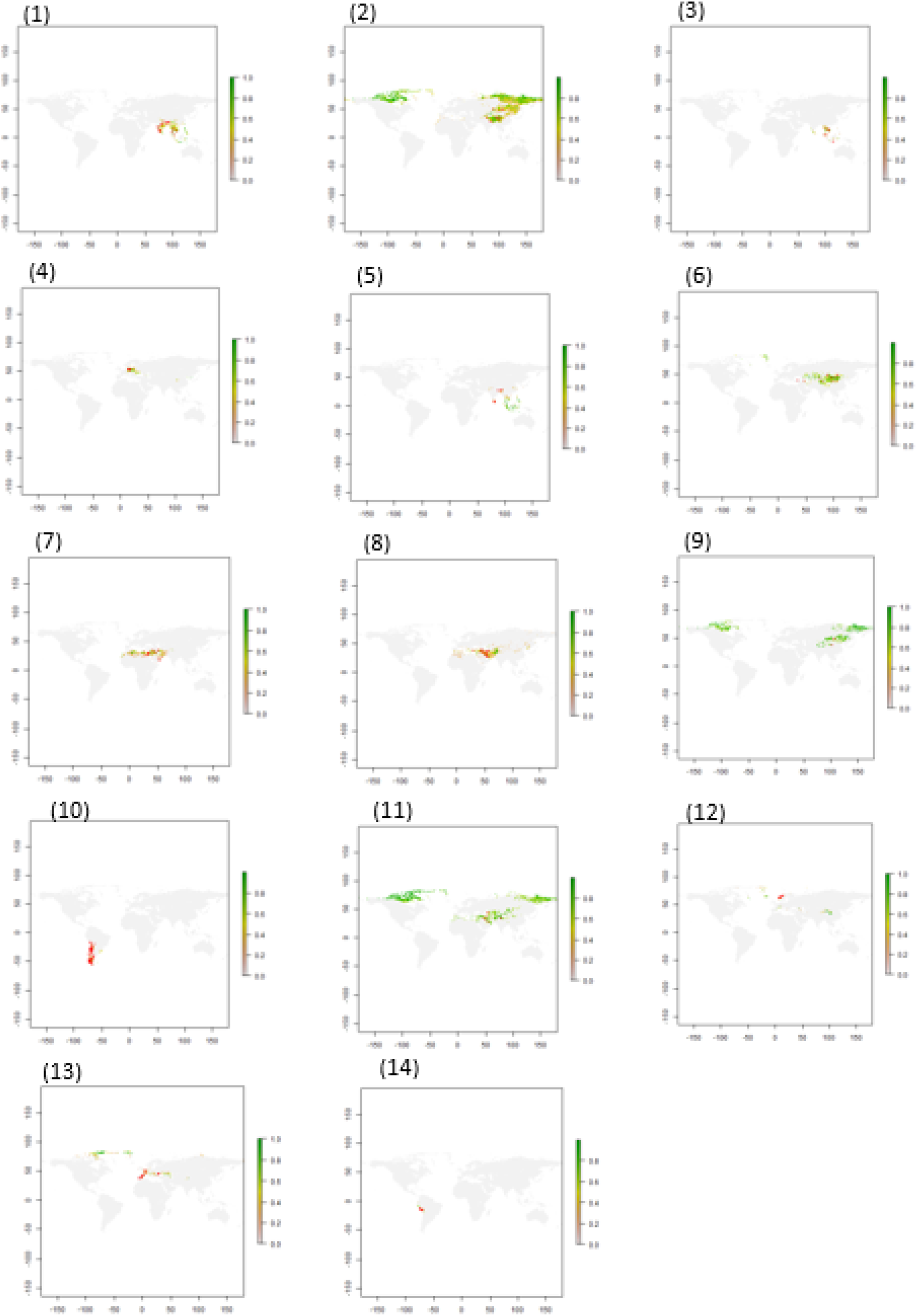
Restricted modelled habitat projections for all domesticated Ungulates. The red dots represent the occurrence locations. The number represent the following species: (1) *Bos frontalis gaurus*, (2) *Bos grunniens mutus*, (3) *Bos javanicus*, (4) *Bos taurus primigenius*, (5) *Bubalus bubalis arnee*, (6) *Camelus bactrianus*, (7) *Camelus dromedarius*, (8) *Capra hircus aegagrus*, (9) *Equus przewalskii*, (10) *Lama glama guanicoe*, (11) *Ovis aries orientalis*, (12) *Rangifer tarandus*, (13) *Sus scrofa*, (14) *Vicagna vicugna*.

All the separate ENMs for the wild and domesticated Ungulates can be downloaded and viewed per species at our GitHub repository (https://github.com/naturalis/trait-geo-diverse-ungulates/tree/master/results).

In the following paragraph we describe the combined results of all ENMs, see Supplementary Note 1 for a summary per model (AUC values, number of occurrence points, variable importance in the models). Figure 2 shows the mean variable importance (0 to 10) for all the separate models per species and the amount of times the variable is used in the models (0 to 160). The ENMs could only be constructed with variables that were not correlated, within the group of correlated variables one is randomly chosen to be used in the model. As visible in the figure 2 most climate variables were highly correlated and therefore not used as often as the topographic variables and soil characteristics whom were less correlated to one another. The mean importance values show which raster layers had the highest importance in the habitat projections. The results in figure 2 suggest that the spatial distribution of Ungulates is highly dependent on climate variables such as climaticMoistureIndex, the growingDegDay5 and aridityindexThornthwaite. Topograpic characteristics played a small role in the spatial distribution with slope and aspect reaching a 31th and 40th place respectively. The soil characteristics also had a relatively small impact on the models with pHCaCL performing best at a 24th place.

**Fig. 2.**
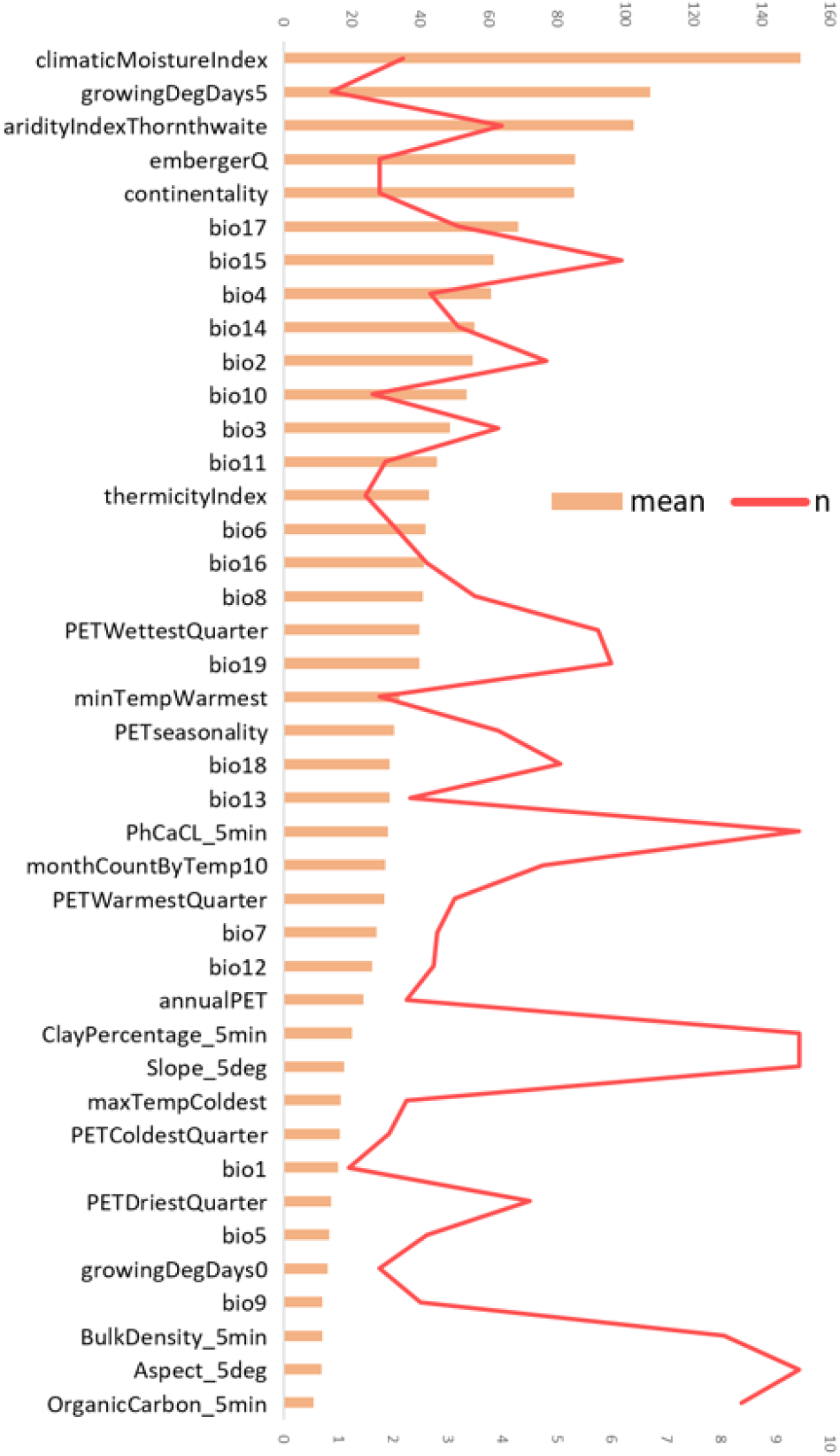
Mean variable importance in the environmental niche models is shown in orange (ranging from 0 to 10), the amount of times that the variable is used in the models is shown in red (ranging from 0 to 160).

### J. Overlap in niche trait space based on modelled habitat projections

The mean niche traits per species can be found on our GitHub repository. The niche traits were used to calculate Gower’s distance in niche trait space between the different species. Figure 3 shows the pairwise patristic distance between domesticated Ungulates and domesticated Ungulates (red line) and the normal distribution of distance (blue bars) between domesticated Ungulates and random sampled wild Ungulates. The mean distance in niche trait space between domesticated Ungulates is significantly larger (2 standard deviations) than the mean distance between domesticated Ungulates and wild Ungulates. Figure 4 shows the distances in niche trait space plotted on a distance dendrogram. The dendogram is divided into 31 clusters and the numbers represent the domesticated species, see Supplementary note 2 for the complete list of species per cluster. The domesticated species are mainly clustered on the right side of the dendrogram, with some species clustering together while others are alone in a separate cluster. Resulting are three clusters in which domesticated species cluster together. The first cluster (0) include the the *Capra hircus aegagrus* (8) and *Camelus bactrianus* (6). The second cluster (h) include the *Bos frontalis gaurus* (1) and the *Bos javanicus* (3). The third and last cluster (l) include the *Equus przewalskii* (9), *Bos grunniens mutus* (2) and the *Ovis aries orientalis* (11). All clusters differentiate from the other clusters based on a few niche traits. Cluster (o) is best explained by climate variables e.g. PETseasonality, PhCaCL and bio7. The cluster is worst explained by bio5, aspect and PETDriestQuarter. Cluster (h) is best explained by bio13, bio18 and bio16. The cluster is worst explained by PETWarmestQuarter, slope and aspect. Cluster (l) is best explained by bio11, maxTempColdest and bio6, and least explained by slope, aspect and aridityIndex-Thornthwaite. The mean niche traits per species were in a generalized linear prediction model to predicts whether a species is domesticated or not. The species prediction model yielded the best results with an AIC of 67.33 when based only based on the bio4 variable. Some wild Ungulates have similar niche traits to the domesticated Ungulates. The species that came closest to the niche traits of domesticated species based on modelled habitat projections were:

**Fig. 3.**
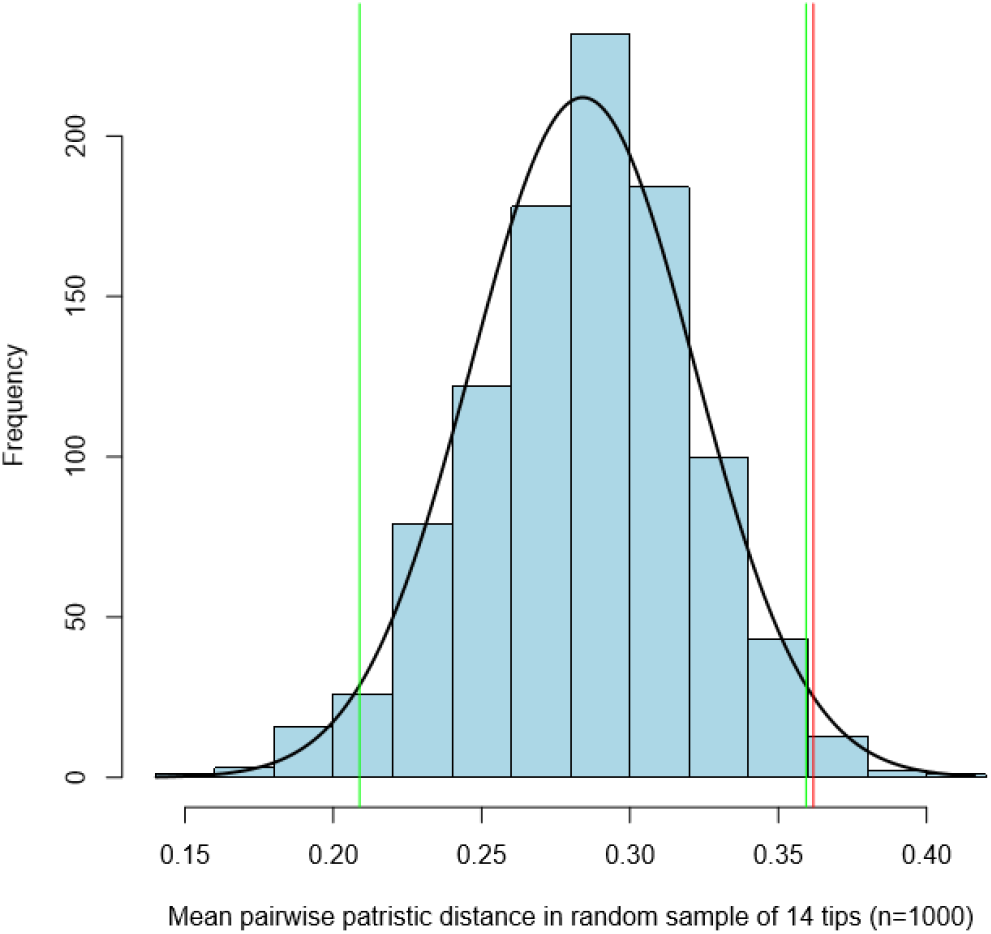
Mean pairwise patristic distance between domesticated Ungulates and domesticated Ungulates (red) and the normal distribution (blue bars) between domesticated Ungulates and random sampled wild Ungulates. The green lines represent 2 standard deviations. Based on the mean normalized niche traits derived from modelled habitat projections.

**Fig. 4.**
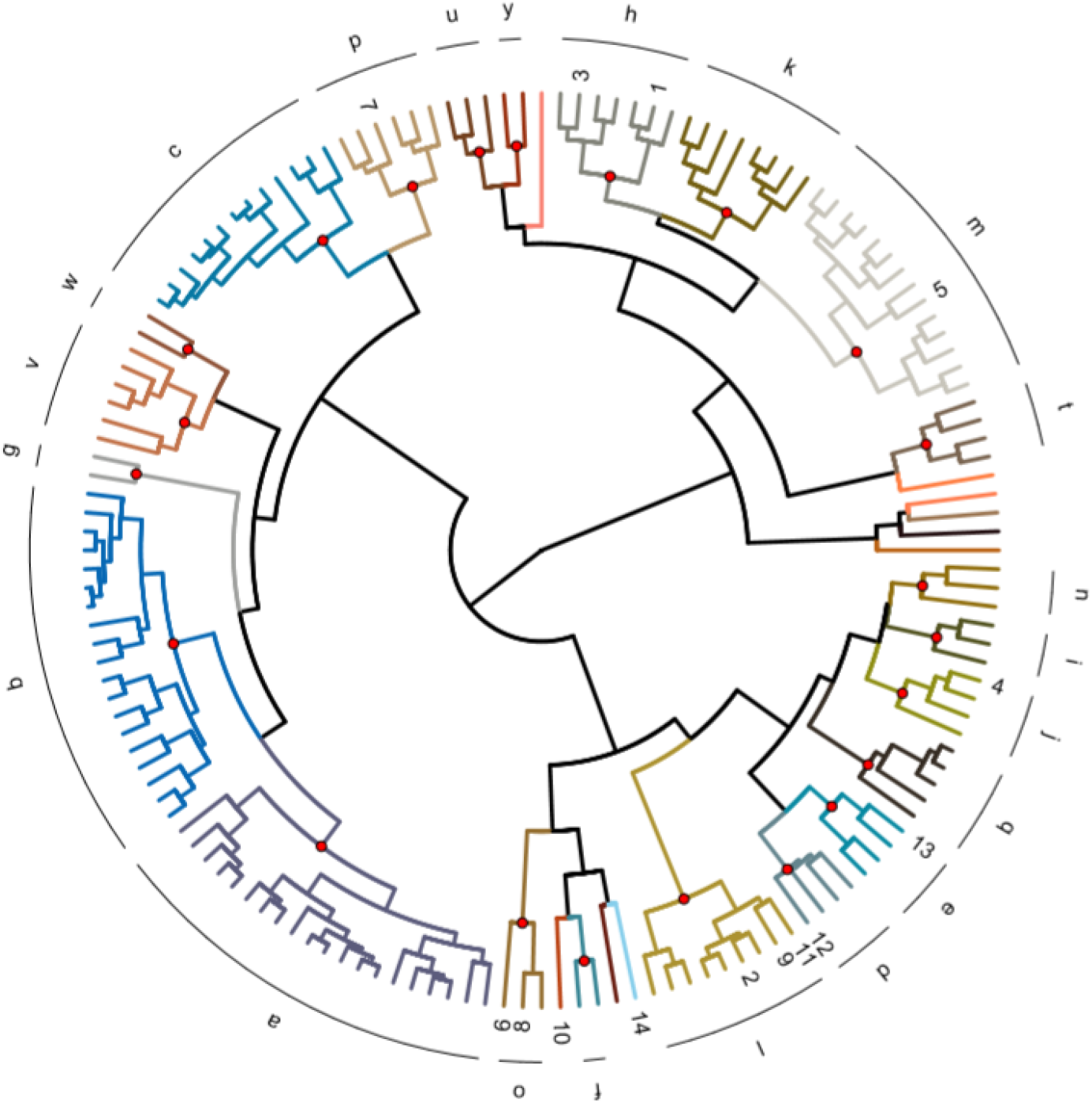
The dendrogram based on modelled habitat projections shows the distances in niche traits between the Ungulates measure with Gower’s D. The species are divided into 31 clusters and the numbers represent the domesticated species. (1) *Bos frontalis gaurus*, (2) *Bos grunniens mutus*, (3) *Bos javanicus*, (4) *Bos taurus primigenius*, (5) *Bubalus bubalis arnee*, (6) *Camelus bactrianus*, (7) *Camelus dromedarius*, (8) *Capra hircus aegagrus*, (9) *Equus przewalskii*, (10) *Lama glama guanicoe*, (11) *Ovis aries orientalis*, (12) *Rangifer tarandus*, (13) *Sus scrofa*, (14) *Vicagna vicugna*.

1. *Capra pyrenaica* (cluster j). Clustered with *Bos taurus primigenius*
2. *Capreolus pygargus* (cluster e). Clustered with *Sus scrofa*.
3. *Eudorcas thomsonii* (cluster v). Clustered without domesticates.
4. *Hemitragus jemlahicus* (cluster k). Clustered without domesticates.
5. *Odocoileus hemionus* (cluster e). Clustered without domesticates.
6. *Odocoileus virginianus* (cluster h). Clustered without domesticates
7. *Ovis ammon* (cluster j). Clustered with *Equus przewalskii*, *Bos grunniens mutus* and *Ovis aries orientalis*.
8. *Pelea capreolus* (cluster v). Clustered without domesticates.

### K. Overlap in niche space derived from modelled habitat projections

Figure 5 shows the pairwise patristic distance between domesticated Ungulates and domesticated Ungulates and wild Ungulates. The distances are based on the overlap in suitable areas measured with Schoener’s D based on the modelled habitat projections. It is visible in figure 5 that the distance between domesticated Ungulates (red line) is smaller (not significant) than the average distance between domesticated Ungulates and wild Ungulates.

**Fig. 5.**
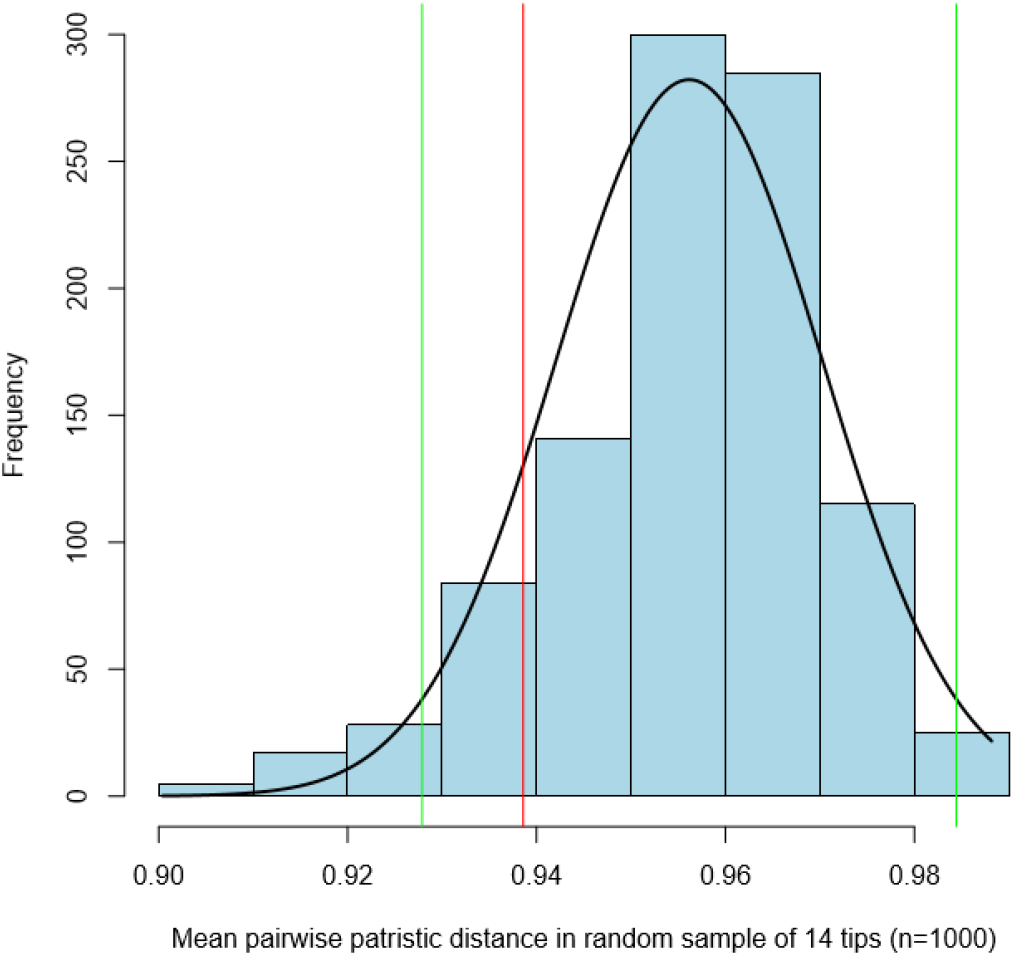
Mean pairwise patristic distance between domesticated Ungulates and domesticated Ungulates (red) and the normal distribution (blue bars) between domesticated Ungulates and random sampled wild Ungulates. The green lines represent 2 standard deviations. Based on niche overlap derived from modelled habitat projections.

The distances in niche overlap are plotted on the dendrogram in figure 6. The dendrogram is divided into 29 clusters, see Supplementary note 3 for the complete list of species per cluster. The domesticated species are dispersed over the dendrogram, some clustering together while others do not. Resulting are four clusters containing multiple domesticated Ungulates. The first cluster (f) contain the *Bos grunniens mutus* (2), the *Ovis aries orientalis* (11), the *Camelus dromedarius* (7) and the *Equus przewalskii* (9). The second cluster (c) contain the *Bos taurus primigenius* (4) and the *Rangifer tarandus* (12). The third cluster contain (b) the *Camelus bactrianus* (6) and the *Capra hircus aegagrus* (8). The fourth and last cluster (g) contain the *Bubalus bubalis arnee* (5), the *Bos frontalis gaurus* (1) and the *Bos javanicus* (3). However, these clusters are not in a near proximity of each other.

**Fig. 6.**
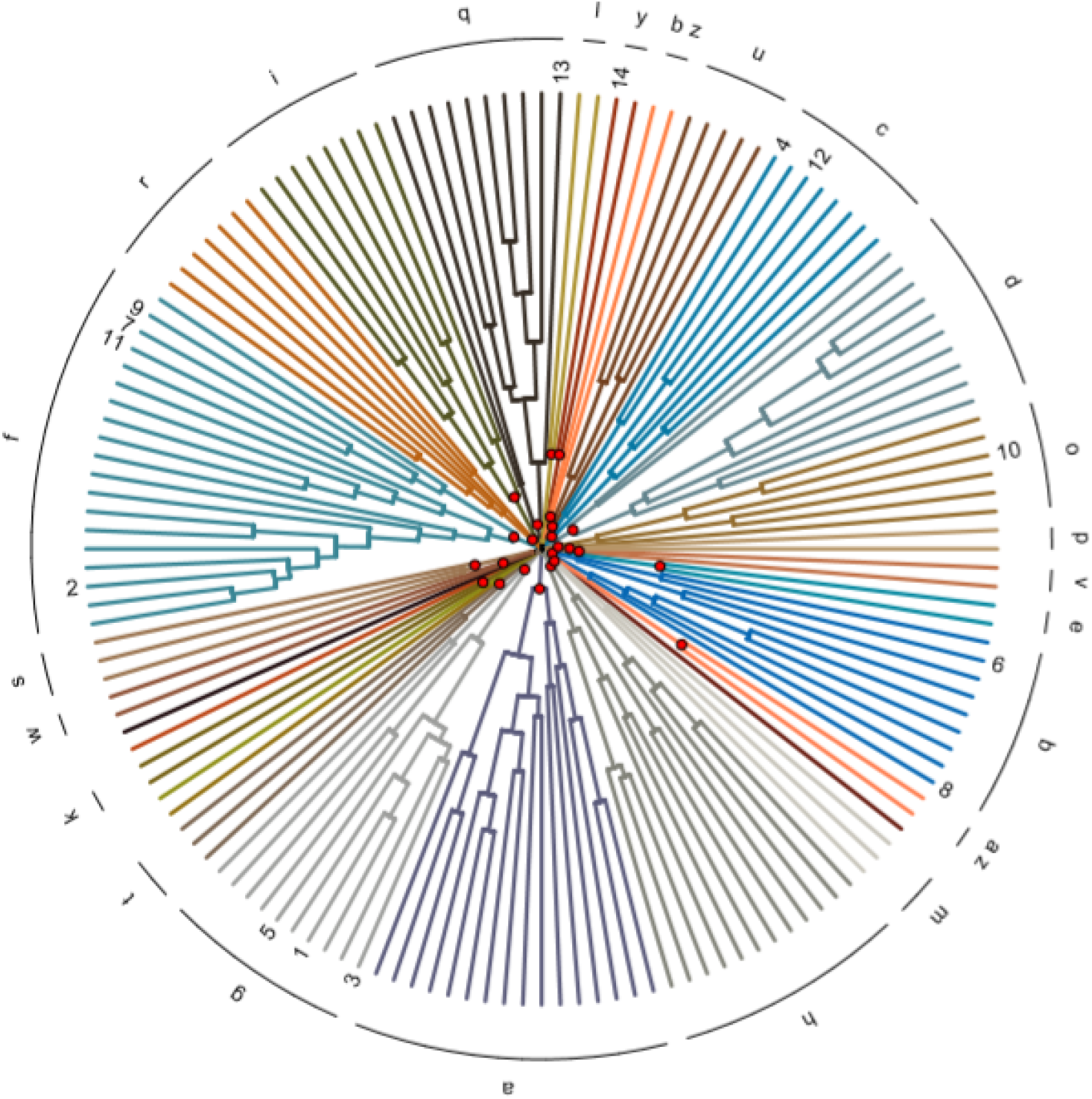
The dendrogram based on modelled habitat projections shows the distances in suitable areas between the Ungulates measure with Schoener’s D. The species are divided into 29 clusters and the numbers represent the domesticated species. (1) *Bos frontalis gaurus*, (2) *Bos grunniens mutus*, (3) *Bos javanicus*, (4) *Bos taurus primigenius*, (5) *Bubalus bubalis arnee*, (6) *Camelus bactrianus*, (7) *Camelus dromedarius*, (8) *Capra hircus aegagrus*, (9) *Equus przewalskii*, (10) *Lama glama guanicoe*, (11) *Ovis aries orientalis*, (12) *Rangifer tarandus*, (13) *Sus scrofa*, (14) *Vicagna vicugna*.

### L. Overlap in niche trait space derived from raw occurrence data

The mean niche traits per species obtained from the raw occurrence data can be viewed and downloaded from our GitHub repository. Figure 7 shows the pairwise patristic mean distance between domesticated Ungulates and domesticated Ungulates (red line) and the distance between domesticated Ungulates and wild Ungulates (blue bars). It is visible that the distance between domesticates is larger (but not significant) than the distance between domesticated Ungulates and wild Ungulates.

**Fig. 7.**
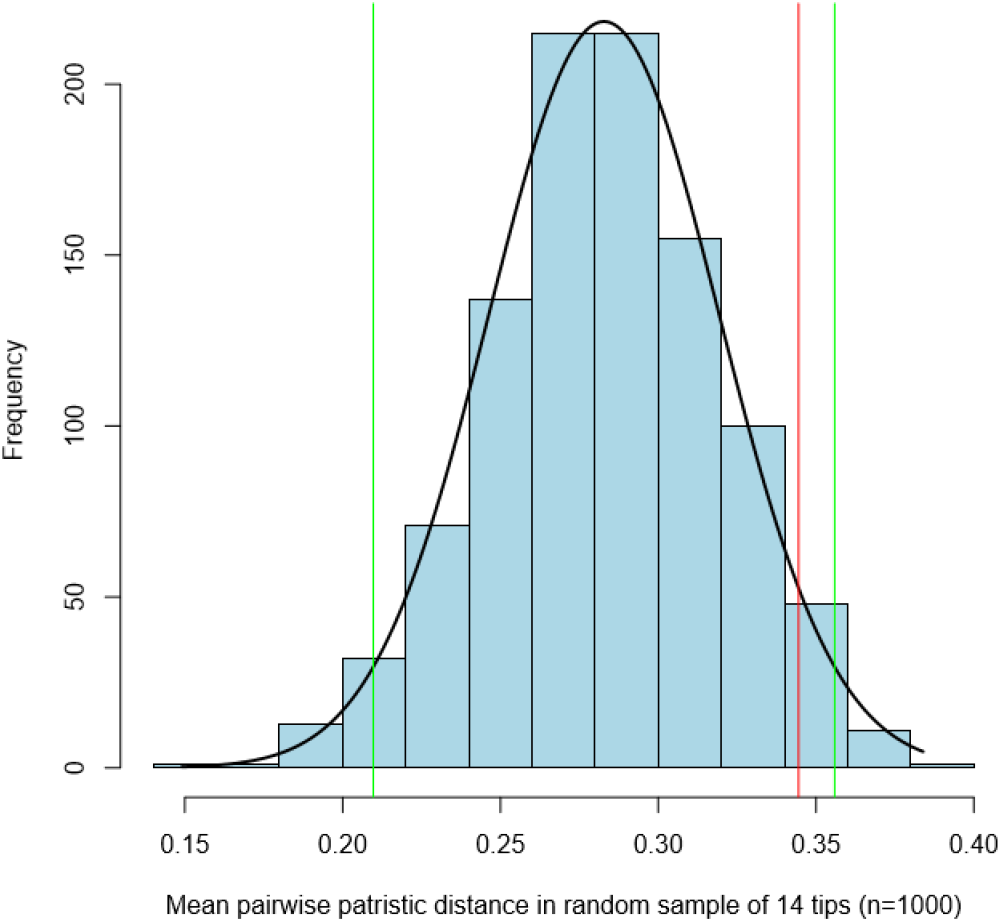
Mean pairwise patristic distance between domesticated Ungulates and domesticated Ungulates (red) and the normal distribution (blue bars) between domesticated Ungulates and random sampled wild Ungulates. The distances in niche traits are based on the raw occurrence data. The green lines represent 2 standard deviations. Based on raw occurrence data

The distances between all Ungulates were plotted on a dendrogram in figure 8. The dendrogram was divided into 31 clusters. It is visible that the species mainly cluster on the top, left and right part leaving the bottom part of the dendrogram without domesticates. Resulting in four cluster with multiple domesticates. The first cluster (m) contains the *Bos javanicus* (3) and the *Bubalus bubalis arnee* (5). The second cluster (j) contains the *Bos taurus primigenius* (4) and the *Sus scrofa* (13). The third cluster (o) contains the *Ovis aries orientalis* (11) and the *Camelus bactrianus* (6). The fourth and last cluster (l) contains the *Bos grunniens mutus* (2) and the *Equus przewalskii* (9). Three of the four clusters are located on the top right side of the dendrogram with the exception of cluster (m) on the left side.

**Fig. 8.**
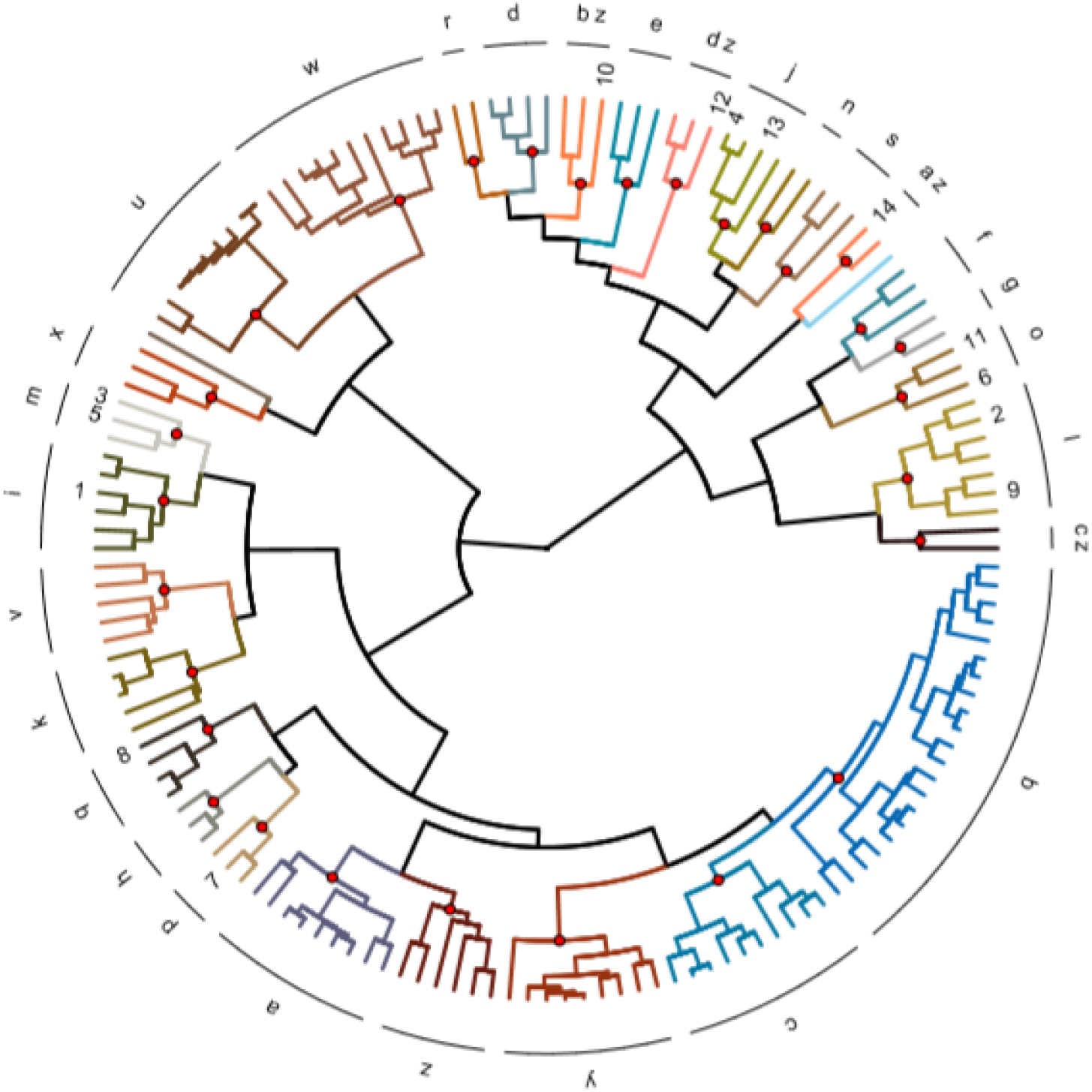
The dendrogram shows the distances in niche trait space between the Ungulates measure with Gower’s D based on raw occurrence data. The species are divided into 31 clusters and the numbers represent the domesticated species.(1) *Bos frontalis gaurus*, (2) *Bos grunniens mutus*, (3) *Bos javanicus*, (4) *Bos taurus primigenius*, (5) *Bubalus bubalis arnee*, (6) *Camelus bactrianus*, (7) *Camelus dromedarius*, (8) *Capra hircus aegagrus*, (9) *Equus przewalskii*, (10) *Lama glama guanicoe*, (11) *Ovis aries orientalis*, (12) *Rangifer tarandus*, (13) *Sus scrofa*, (14) *Vicagna vicugna*.

The clusters deviate from another based on a few important niche traits. Cluster (m) is best explained by climate variables e.g. bio13, bio16 and bio18. The cluster is worst explained by BulkDensity, Aspect and Slope. Cluster (j) on the other hand is best explained by bio3, PETColdestQuarter and bio15 and worst explained by climaticMoistureIndex, aspect and PETWettestQuarter. Cluster (o) is best explained by PETseasonality, bio13 and bio12. The cluster is worst explained by soil and topography characteristics e.g. bio5, aspect and ClayPercentage. Cluster (l) is best explained by monthCountByTemp10, thermicityIndex and minTempWarmest. The cluster is worst explained by Aspect, Carbon and BulkDensity.

The mean niche traits per species were also used in a generalized linear prediction model that predicts whether a species is domesticated or not. The generalized linear prediction model that yielded the lowest AIC of 61.97 was best explained by bio3, OrganicCarbon, bio2, bio17 and BulkDensity.

When applying this model to the whole dataset 8 wild species have similar traits to domesticated species.

1. *Ammotragus lervia* (cluster f). Clustered without other domesticates.
2. *Damaliscus pygargus* (cluster z). Clustered without other domesticates.
3. *Muntiacus muntjak* (cluster m). Clustered with (Bos javanicus and *Bubalus bubalis arnee*.
4. *Ovis ammon* (cluster l). Clustered with *Bos grunniens mutus* and *Equus przewalskii*.
5. *Pelea capreolus* (cluster z). Clustered without other domesticates.
6. *Potamochoerus larvatus* (cluster a). Clustered without other domesticates.
7. *Raphicerus melanotis* (cluster z). Clustered without other domesticates.
8. *Rusa unicolor* (cluster i). Clustered with *Bos frontalis gaurus*.

## Discussion

### M. Ecological niche models

The modelled habitat projections in figure 1 show that some models are likely to be overfitted on one variable. This is very clear for the *Bos grunniens mutus* (2), *Ovis aries orientalis* (11) and *Equus przewalskii* (9) where suitable areas include the most northern areas on earth (Canada and Greenland). The aforementioned species are unlikely to sustain these harsh environments and therefore the models are not accurate enough for these species. The overfitting is likely due to the importance of a few variables such as slope and bio14. These variables have similar values in areas around Mongolia, Russia and Tibet as areas in and around Canada and Greenland. The model does not take variables such as minimum and maximum temperatures into account because this was not of importance in the model environment in which the models were trained. When projecting the suitable habitat areas on the whole earth these variables are of significant importance.

The problem might be solved by applying different restricted areas. For this research we restricted the predictions by Holt’s zoogeographical regions which divide the earth in 11 large realms (51). Unrealistic modelled habitat projections only seem to occur for the Ungulates with occurrence points that are inside the palearctic realm (including Europe, Russia, Asia, Greenland and Canada). Future work could focus on the use of different or stricter realms when restricting habitat projection models.

Another limit of the use of ENM is the amount of input data needed to accurately describe biomes and interactions between environmental variables. We used Hutchinsons description of a niche as a n-dimensional hypervolume where environmental variables explain the dimensions. In this research we used n=41 dimensions which is not enough to describe complex systems such as niches. The probability is high that our model lacks important climate, topography and soil variables that explain the distribution of species better. Also important interactions between variables could be missed. Further research is needed on the availability, use and development of variables that explain ecological niches.

### N. Comparison of niche overlap methods

The pairwise patristic distances calculated with three different methods show contrasting results. The pairwise differences calculated with (i) overlap in niche trait space derived from modelled habitat projections and (ii) niche trait space overlap based on raw occurrence data measured both measured with Gower’s distance show that the average distance between domesticates is larger than the distance between domesticated and wild Ungulates. This suggests that domesticates are a very versatile group with little niche similarities. The dendrograms in figure 4 and 8 show that the domesticated species are widely dispersed and it is not possible to cluster the domesticated species together. However, it is possible to cluster types of domesticates together.

The pairwise patristic distance calculated with the niche space based on modelled habitat projections measured with Schoener’s D shows that the average distances between domesticates is smaller than the average distances between domesticated and wild Ungulates. It might be possible that wide niches are better captured by niche space because in the calculation of niche traits the niche traits are normalized and averaged removing the variety in the data set. Further research into the effects over normalization and averaging environmental variables is needed.

The dendrograms in figure 4, 6 and 8 do not show one clear cluster with all domesticated species in it but rather a few clusters of domesticates. When comparing the dendrograms based on the three used methods some similarities and differences can be found. The *Bubalus bubalis arnee*, *Bos javanicus* and *Bos frontalis gaurus* are always in the same cluster or on the same major tree branch. This clustering seems reasonable since they all reside in the same areas (see figure 1.) The MaxEnt projections show an accurate representation of these species without overfitting and projections in unexpected areas. The differences in dendrograms are mainly visible for the species that suffered overfitting problems in the MaxEnt projections *Sus scrofa*, *Bos grunniens mutus*, *Ovis aries orientalis* and *Equus przewalskii*.

Overall we can conclude that the problems related to the ENMs make the results unreliable, suggesting that the dendrograms based on modelled habitat projections are less reliable than the dendrogram based on raw occurrence points. The modelled habitat projections project suitable areas in places that are uninhabitable for the species. Therefore the clustering based on the raw occurrence points seems the most accurate. Our results based on environmental traits obtained from raw occurrence points show that the variation between domesticated species is to great to capture a set of common environmental traits for all domesticated Ungulates.

The same conclusion was shown by the generalized linear prediction model which was used to assess which variables are best to predict domestication and whether some wild Ungulates resemble the domesticates based on niche traits. The wild species that resembled the domesticated species were sometimes in the same cluster as the domesticates but they were also in clusters without domesticates.

Our results shows that domesticates are a very versatile group with different purposes for domestication. Some species are mainly used for transportation, or for farming (pulling plows) other species are mainly used for the production of food or a combination of purposes. The variety within the domesticated Ungulate group was too big to capture similar niche traits hence further research in the use of other traits is needed. Only 41 environmental traits were tested and there is still a whole range of likely variables that need to be tested.

### O. Evaluation of the framework

To calculate the overlap in niche trait space and niche space per species we constructed a framework that automatically construct ENMs for a given number of species. During the construction of the framework some assumption were made. The first assumption we made was that the niches of the wild ancestors of domesticated Ungulates are the same as the niches of the domesticated Ungulates right before they were domesticated. It might be possible that small changes in their niche preferences developed before they were domesticated. Therefore using wild ancestors of domesticates as a proxy of domesticated species comes with a risk. On the other hand there is not enough occurrence data available of domesticated species before they were bread in captivity to calculate niche trait space and niche space for.

The second assumption involves the comparison of occurrence points before 1950 with climatic data after 1950. In order to obtain enough occurrence points for the wild ancestors of domesticated Ungulates we had to select occurrence points that were measured before 1950. For example the *Bos taurus primigenius*, the wild ancestor of cattle, went extinct around 1627 (65) and therefore older measurements had to be used. The use of older measurements leads to problems related to the environmental data. We extracted climate variables from Bioclim and ENVIREM datasets which are based on monthly remote sensing data between 1950 and 2000 (32), the topographic variables are less likely to change greatly during these relatively short time intervals. For this research we assumed with caution that using the climatic dataset based on our current climate, is enough to account for the climate variability during the course of the late Holocene. Connoly et al., (2012) (42) made a similar assumption since detailed climate variables are only available for large time steps such as mid-holocene and early-holocene with a lower spatial resolution. However we are aware that climatic variability was present during the course of the late Holocene which has been shown in ice cores from Greenland and Antarctic (66).

The third assumption that we made is that mean environmental variables e.g niche traits can capture the niche of a species. When normalizing and averaging environmental variables the whole variety in the data set is lost. For example it might be possible that a species functions well in areas with a rainfall between 200 and 500 mm per year, in our model it seems like the species only lives in areas with 350 mm rainfall per year. For further research it might be interesting to take the niche breadth of a species into account. Some species function well in a whole range of climate, topographic and soil varieties and might therefore be more similar that the averages suggest.

## Conclusion

The differentiation between domesticated and wild Ungulates is most accurate when using the direct environmental variables extracted from the raw occurrence points. The environmental niche models overfit their projections on a few variables that were of importance in the model environment. Outside of the model environment other environmental variables limit the distribution of species which the model does not deem important. Therefore more research into the construction of accurate ENMs is needed.

The dendrogram based on the raw occurrence points shows that some domesticated species cluster together while others do not. The variety within the domesticate Ungulates group is to large to differentiate between them and wild Ungulates based on the 41 niche traits used in this research. Further research into the use of additional niche traits is needed.

## ACKNOWLEDGEMENTS

We would like to express our appreciation to Dr. Niels Raes for his advice and technical support on the implementation of Ecological Niche Modelling. We would also like to thank Tom van Dooren for his advise on the use of the Outlying Mean Index. Finally, we wish to thank the research group *Understanding Evolution*, part of Naturalis biodiversity center, for hosting us during the course of this research.

## Supplementary Note 1 Summary environmental niche models

**Table 4.**
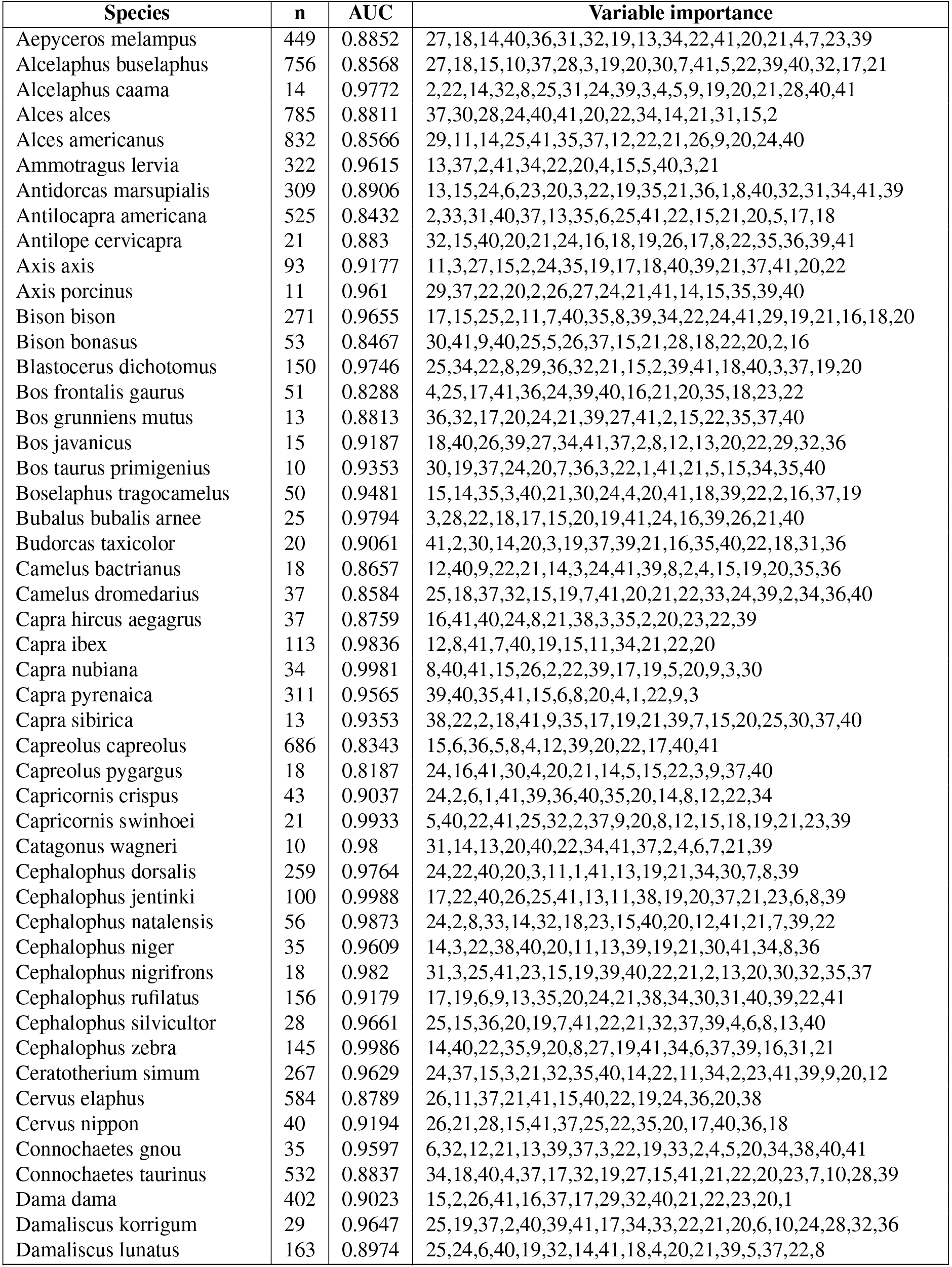
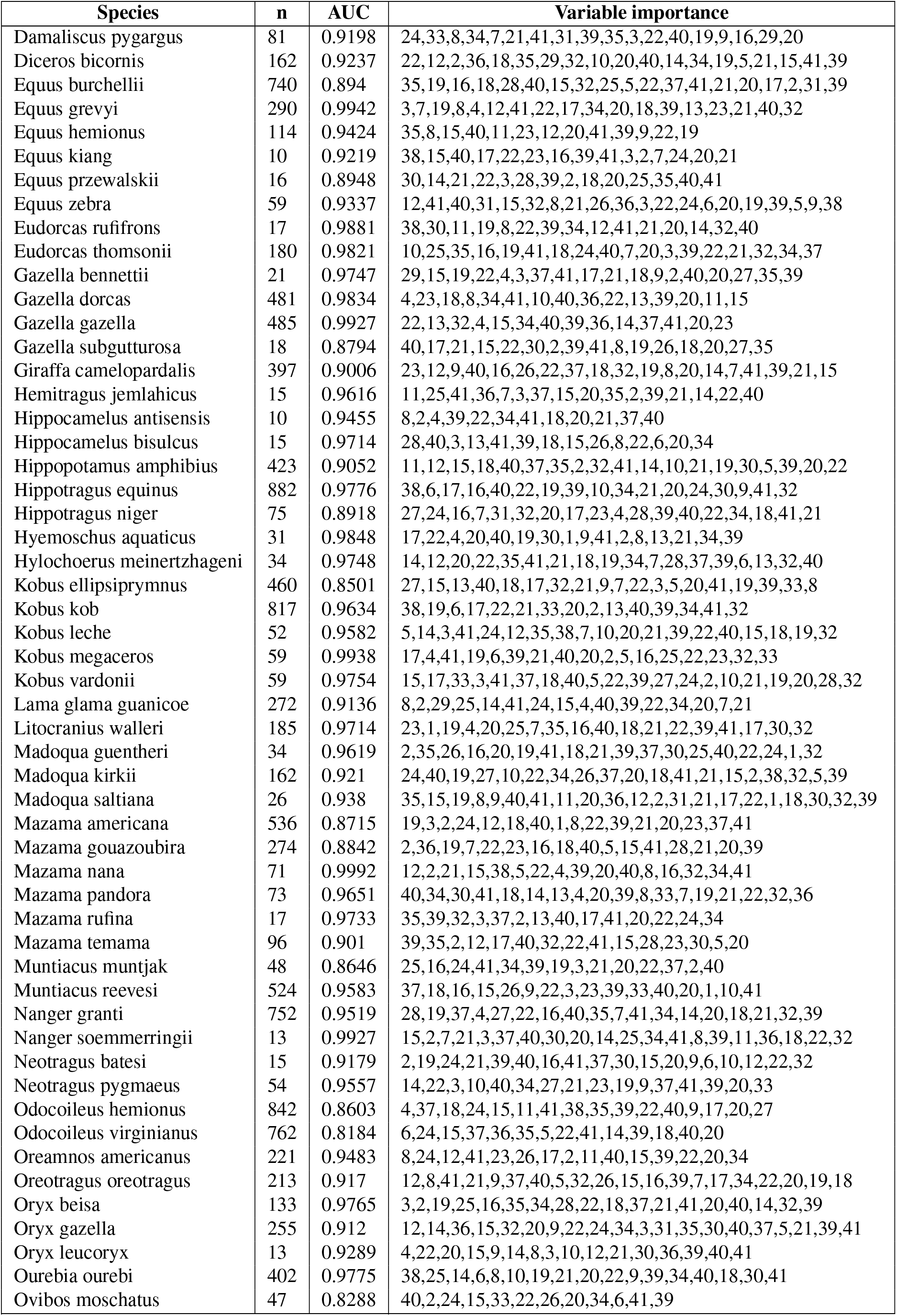
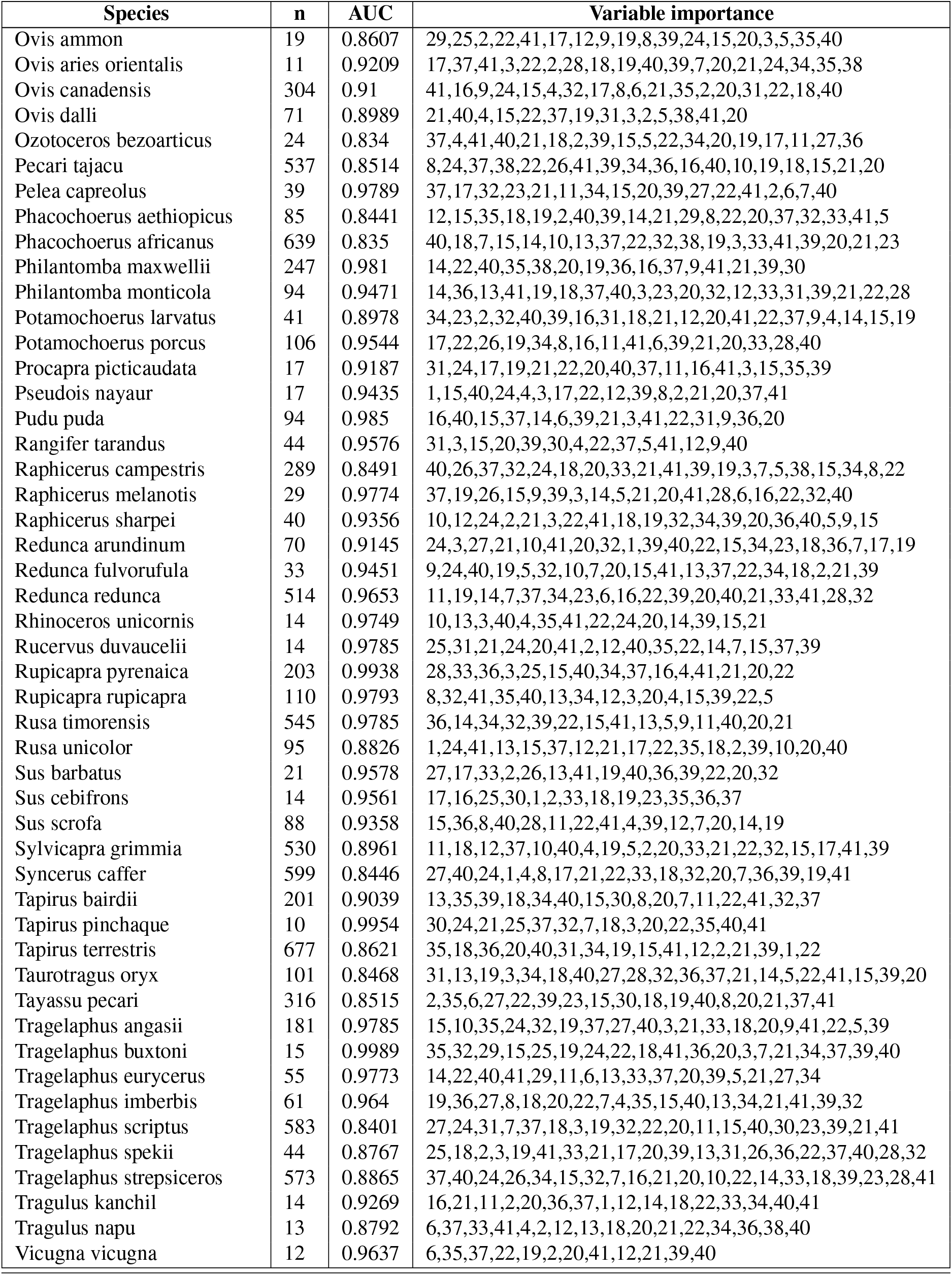
Summary of the environmental niche models with the amount of occurrence points AUC value and variables used in the model in a descending list from most important to least important. Where 1 = bio1, 2 = bio2, 3 = bio3, 4 = bio4, 5 = bio5, 6 = bio6, 7 = bio7, 8 = bio8, 9 = bio9, 10 = bio10, 11 = bio11, 12 = bio12, 13 = bio13, 14 = bio14, 15 = bio15, 16 = bio16, 17 = bio17, 18 = bio18, 19 = bio19, 20 = Aspect, 21 = BulkDensity, 22 = ClayPercentage, 23 = annualPET, 24 = aridityIndexThornthwaite, 25 = climaticMoistureIndex, 26 = continentality, 27 = embergerQ, 28 = growingDegDays0, 29 = growingDegDays5, 30 = maxTempColdest, 31 = minTempWarmest, 32 = monthCountByTemp10, 33 = PETColdestQuarter, 34 = PETDriestQuarter, 35 = PETseasonality, 36 = PETWarmestQuarter, 37 = PETWettestQuarter, 38 = thermicityIndex, 39 = OrganicCarbon, 40 = PhCaCL and 41 = slope

## Supplementary Note 2: Clusters of species in niche trait space based on modelled habitat projections

**Table 5.**
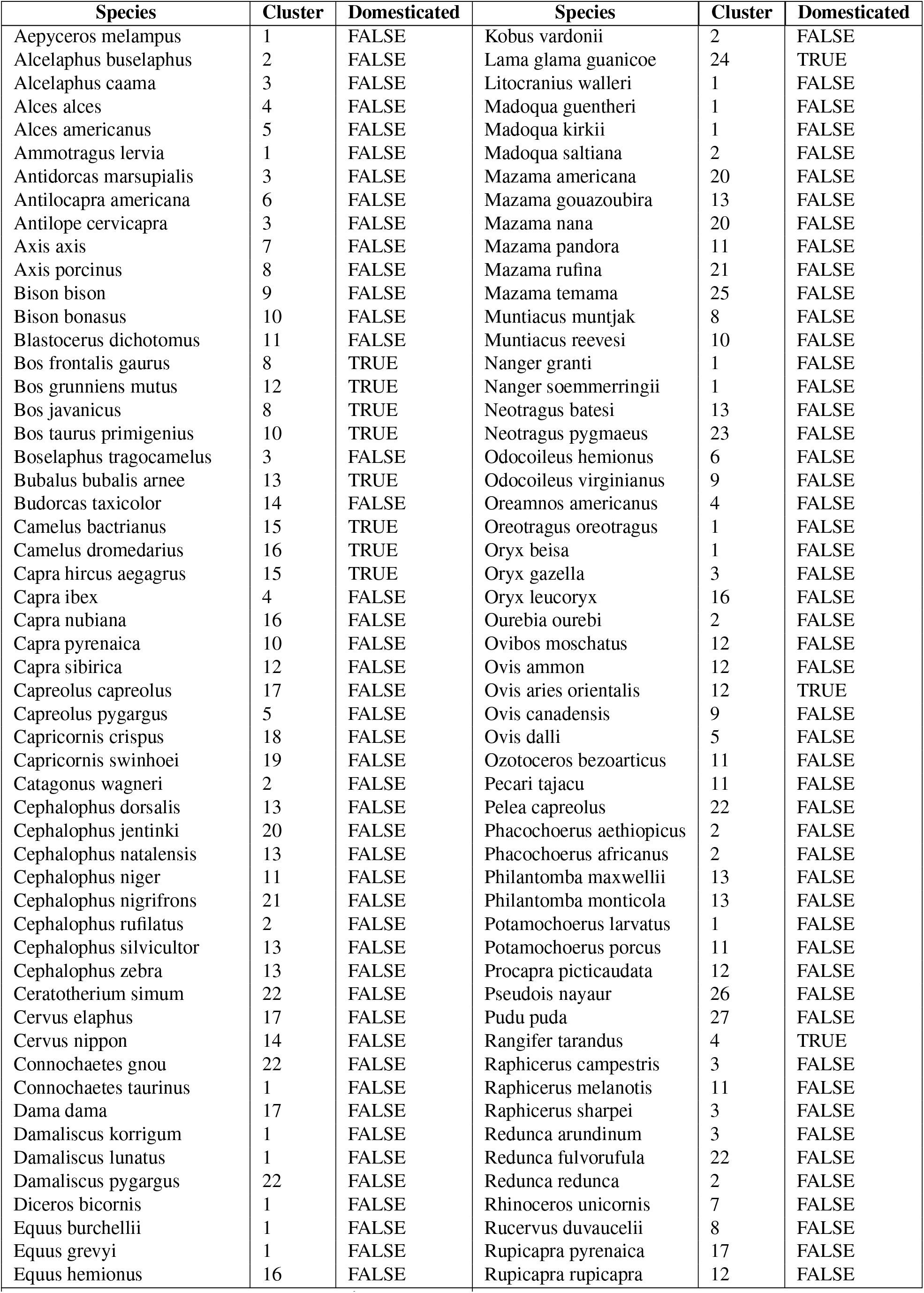
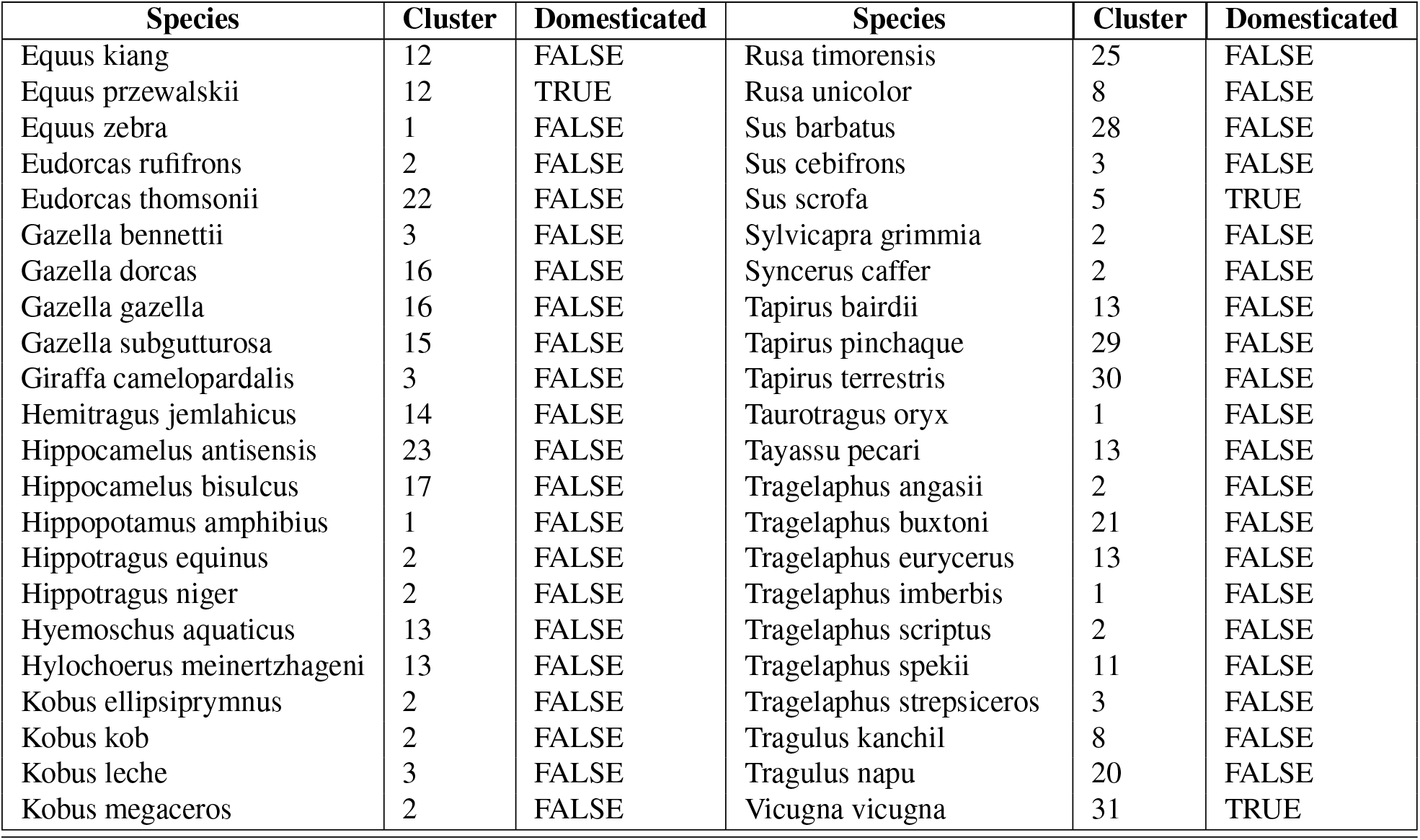
Species per cluster based on niche traits space with modelled habitat projections

## Supplementary Note 3: Clusters of species in niche space based on modelled habitat projections

**Table 6.**
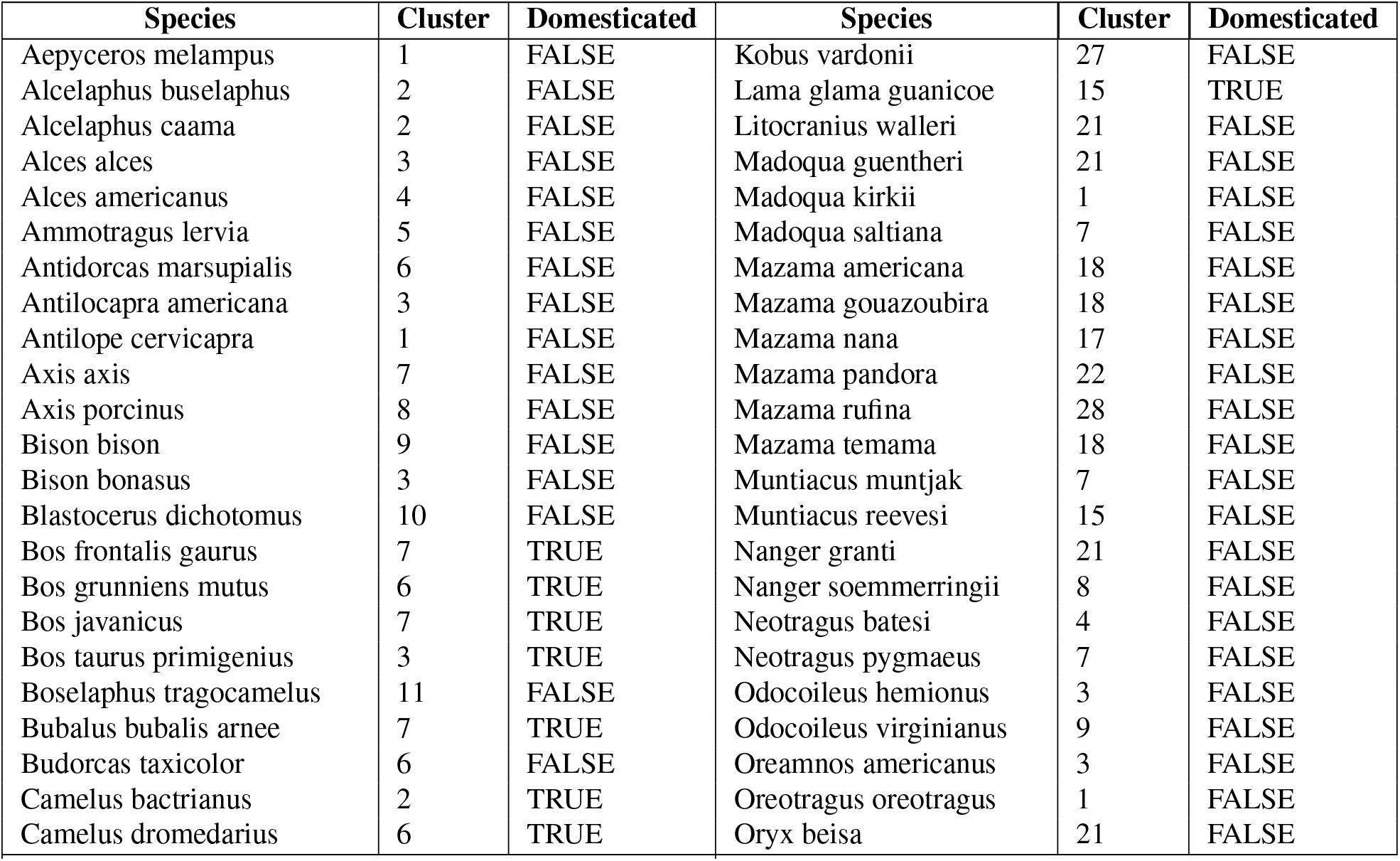
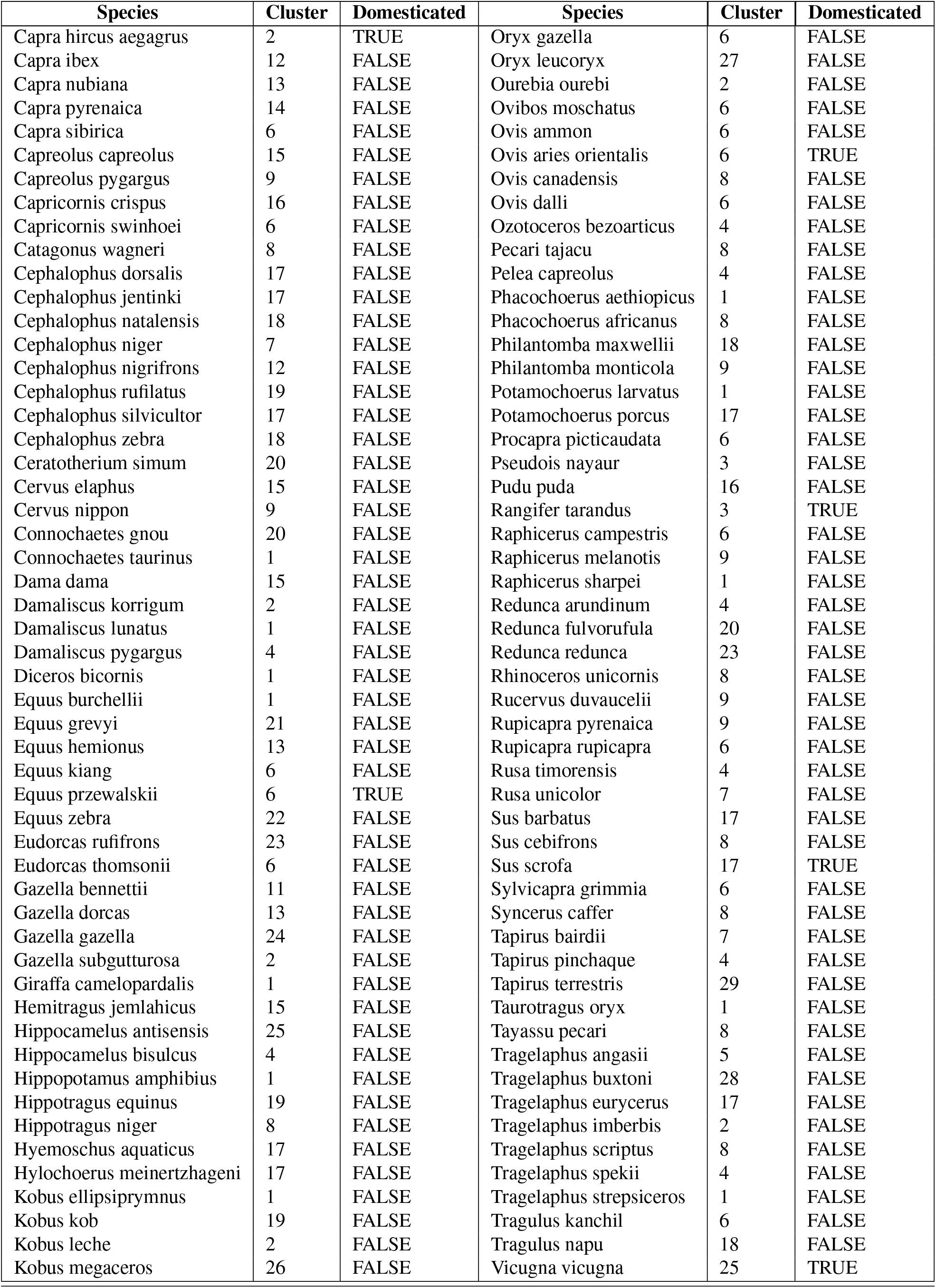
Clusters of species in niche space based on modelled habitat projections

## Supplementary Note 4: Clusters of species based on raw occurrences

**Table 7.** Species per cluster based on raw occurrences

| Species | Cluster | Domesticated | Species | Cluster | Domesticated |
| --- | --- | --- | --- | --- | --- |
| Aepyceros melampus | 1 | FALSE | Kobus vardonii | 2 | FALSE |
| Alcelaphus buselaphus | 2 | FALSE | Lama glama guanicoe | 28 | TRUE |
| Alcelaphus caama | 3 | FALSE | Litocranius walleri | 2 | FALSE |
| Alces alces | 4 | FALSE | Madoqua guentheri | 2 | FALSE |
| Alces americanus | 5 | FALSE | Madoqua kirkii | 1 | FALSE |
| Ammotragus lervia | 6 | FALSE | Madoqua saltiana | 2 | FALSE |
| Antidorcas marsupialis | 3 | FALSE | Mazama americana | 11 | FALSE |
| Antilocapra americana | 7 | FALSE | Mazama gouazoubira | 23 | FALSE |
| Antilope cervicapra | 8 | FALSE | Mazama nana | 11 | FALSE |
| Axis axis | 9 | FALSE | Mazama pandora | 22 | FALSE |
| Axis porcinus | 9 | FALSE | Mazama rufina | 24 | FALSE |
| Bison bison | 5 | FALSE | Mazama temama | 23 | FALSE |
| Bison bonasus | 10 | FALSE | Muntiacus muntjak | 13 | FALSE |
| Blastocerus dichotomus | 11 | FALSE | Muntiacus reevesi | 18 | FALSE |
| Bos frontalis gaurus | 9 | TRUE | Nanger granti | 2 | FALSE |
| Bos grunniens mutus | 12 | TRUE | Nanger soemmerringii | 2 | FALSE |
| Bos javanicus | 13 | TRUE | Neotragus batesi | 23 | FALSE |
| Bos taurus primigenius | 10 | TRUE | Neotragus pygmaeus | 21 | FALSE |
| Boselaphus tragocamelus | 8 | FALSE | Odocoileus hemionus | 6 | FALSE |
| Bubalus bubalis arnee | 13 | TRUE | Odocoileus virginianus | 10 | FALSE |
| Budorcas taxicolor | 14 | FALSE | Oreamnos americanus | 5 | FALSE |
| Camelus bactrianus | 15 | TRUE | Oreotragus oreotragus | 1 | FALSE |
| Camelus dromedarius | 16 | TRUE | Oryx beisa | 2 | FALSE |
| Capra hircus aegagrus | 17 | TRUE | Oryx gazella | 3 | FALSE |
| Capra ibex | 18 | FALSE | Oryx leucoryx | 16 | FALSE |
| Capra nubiana | 17 | FALSE | Ourebia ourebi | 25 | FALSE |
| Capra pyrenaica | 6 | FALSE | Ovibos moschatus | 29 | FALSE |
| Capra sibirica | 12 | FALSE | Ovis ammon | 12 | FALSE |
| Capreolus capreolus | 4 | FALSE | Ovis aries orientalis | 15 | TRUE |
| Capreolus pygargus | 19 | FALSE | Ovis canadensis | 7 | FALSE |
| Capricornis crispus | 19 | FALSE | Ovis dalli | 29 | FALSE |
| Capricornis swinhoei | 20 | FALSE | Ozotoceros bezoarticus | 26 | FALSE |
| Catagonus wagneri | 3 | FALSE | Pecari tajacu | 22 | FALSE |
| Cephalophus dorsalis | 21 | FALSE | Pelea capreolus | 26 | FALSE |
| Cephalophus jentinki | 21 | FALSE | Phacochoerus aethiopicus | 2 | FALSE |
| Cephalophus natalensis | 22 | FALSE | Phacochoerus africanus | 2 | FALSE |
| Cephalophus niger | 23 | FALSE | Philantomba maxwellii | 21 | FALSE |
| Cephalophus nigrifrons | 24 | FALSE | Philantomba monticola | 22 | FALSE |
| Cephalophus rufilatus | 25 | FALSE | Potamochoerus larvatus | 1 | FALSE |
| Cephalophus silvicultor | 23 | FALSE | Potamochoerus porcus | 23 | FALSE |
| Cephalophus zebra | 21 | FALSE | Procapra picticaudata | 12 | FALSE |
| Ceratotherium simum | 3 | FALSE | Pseudois nayaur | 12 | FALSE |
| Cervus elaphus | 4 | FALSE | Pudu puda | 28 | FALSE |
| Cervus nippon | 19 | FALSE | Rangifer tarandus | 30 | TRUE |
| Connochaetes gnou | 26 | FALSE | Raphicerus campestris | 3 | FALSE |
| Connochaetes taurinus | 2 | FALSE | Raphicerus melanotis | 26 | FALSE |
| Dama dama | 4 | FALSE | Raphicerus sharpei | 2 | FALSE |
| Damaliscus korrigum | 25 | FALSE | Redunca arundinum | 3 | FALSE |
| Damaliscus lunatus | 2 | FALSE | Redunca fulvorufula | 1 | FALSE |
| Damaliscus pygargus | 26 | FALSE | Redunca redunca | 25 | FALSE |
| Diceros bicornis | 1 | FALSE | Rhinoceros unicornis | 9 | FALSE |
| Equus burchellii | 1 | FALSE | Rucervus duvaucelii | 9 | FALSE |

Table 7 – continued from previous page
| Species | Cluster | Domesticated | Species | Cluster | Domesticated |
| --- | --- | --- | --- | --- | --- |
| *Equus grevyi* | 2 | FALSE | *Rupicapra pyrenaica* | 30 | FALSE |
| *Equus hemionus* | 17 | FALSE | *Rupicapra rupicapra* | 30 | FALSE |
| *Equus kiang* | 12 | FALSE | *Rusa timorensis* | 11 | FALSE |
| *Equus przewalskii* | 12 | TRUE | *Rusa unicolor* | 9 | FALSE |
| *Equus zebra* | 26 | FALSE | *Sus barbatus* | 21 | FALSE |
| *Eudorcas rufifrons* | 25 | FALSE | *Sus cebifrons* | 23 | FALSE |
| *Eudorcas thomsonii* | 1 | FALSE | *Sus scrofa* | 10 | TRUE |
| *Gazella bennettii* | 8 | FALSE | *Sylvicapra grimmia* | 25 | FALSE |
| *Gazella dorcas* | 16 | FALSE | *Syncerus caffer* | 2 | FALSE |
| *Gazella gazella* | 17 | FALSE | *Tapirus bairdii* | 23 | FALSE |
| *Gazella subgutturosa* | 15 | FALSE | *Tapirus pinchaque* | 24 | FALSE |
| *Giraffa camelopardalis* | 2 | FALSE | *Tapirus terrestris* | 11 | FALSE |
| *Hemitragus jemlahicus* | 14 | FALSE | *Taurotragus oryx* | 1 | FALSE |
| *Hippocamelus antisensis* | 27 | FALSE | *Tayassu pecari* | 23 | FALSE |
| *Hippocamelus bisulcus* | 28 | FALSE | *Tragelaphus angasii* | 3 | FALSE |
| *Hippopotamus amphibius* | 2 | FALSE | *Tragelaphus buxtoni* | 31 | FALSE |
| *Hippotragus equinus* | 25 | FALSE | *Tragelaphus eurycerus* | 21 | FALSE |
| *Hippotragus niger* | 2 | FALSE | *Tragelaphus imberbis* | 2 | FALSE |
| *Hyemoschus aquaticus* | 21 | FALSE | *Tragelaphus scriptus* | 2 | FALSE |
| *Hylochoerus meinertzhageni* | 23 | FALSE | *Tragelaphus spekii* | 22 | FALSE |
| *Kobus ellipsiprymnus* | 2 | FALSE | *Tragelaphus strepsiceros* | 3 | FALSE |
| *Kobus kob* | 25 | FALSE | *Tragulus kanchil* | 23 | FALSE |
| *Kobus leche* | 3 | FALSE | *Tragulus napu* | 21 | FALSE |
| *Kobus megaceros* | 25 | FALSE | *Vicugna vicugna* | 27 | TRUE |

## Bibliography

1. Peter Kareiva, Sean Watts, Robert McDonald, and Tim Boucher. Domesticated nature: shaping landscapes and ecosystems for human welfare. Science, 316(5833):1866–1869, 2007.

2. Greger Larson, Dolores R Piperno, Robin G Allaby, Michael D Purugganan, Leif Andersson, Manuel Arroyo-Kalin, Loukas Barton, Cynthia Climer Vigueira, Tim Denham, Keith Dobney, et al. Current perspectives and the future of domestication studies. Proceedings of the National Academy of Sciences, 111(17):6139–6146, 2014.

3. Greger Larson, Elinor K Karlsson, Angela Perri, Matthew T Webster, Simon YW Ho, Joris Peters, Peter W Stahl, Philip J Piper, Frode Lingaas, Merete Fredholm, et al. Rethinking dog domestication by integrating genetics, archeology, and biogeography. Proceedings of the National Academy of Sciences, 109(23):8878–8883, 2012.

4. Anil K Gupta. Origin of agriculture and domestication of plants and animals linked to early holocene climate amelioration. CURRENT SCIENCE-BANGALORE-, 87:54–59, 2004.

5. Bridget Allchin and Frank Raymond Allchin. Origins of a civilization: the prehistory and early archaeology of South Asia. Viking Adult, 1997.

6. Glen Michael MacDonald. Biogeography: space, time and life. Number Sirsi) i9780471241935. 2003.

7. Bruce D Smith and Mark Nesbitt. The emergence of agriculture. Scientific American Library New York, 1995.

8. Bruce D Smith. Documenting plant domestication: The consilience of biological and archaeological approaches. Proceedings of the National Academy of Sciences, 98(4):1324–1326, 2001.

9. Jared M Diamond and Doug Ordunio. Guns, germs, and steel. Books on Tape, 1999.

10. Melinda A Zeder. Domestication and early agriculture in the mediterranean basin: Origins, diffusion, and impact. Proceedings of the national Academy of Sciences, 105(33):11597–11604, 2008.

11. Charles Darwin. The origin of species: By means of natural selection of the preservation of favoured races in the struggle for life. Kennebec Large Print, 2010.

12. Karl Hammer. Das domestikationssyndrom. Die Kulturpflanze, 32(1):11–34, 1984.

13. Jack Rodney Harlan et al. Crops and man. Number Ed. 2. American Society of Agronomy, 1992.

14. Melinda Zeder. The domestication of animals. Journal of Anthropological Research, 68(2): 161–190, 2012.

15. Robert D Holt. Bringing the hutchinsonian niche into the 21st century: ecological and evolutionary perspectives. Proceedings of the National Academy of Sciences, 106(Supplement 2):19659–19665, 2009.

16. G Evelyn Hutchinson. A treatise on limnology. Limnology, 1:243, 1957.

17. Jorge Soberón. Grinnellian and eltonian niches and geographic distributions of species. Ecology letters, 10(12):1115–1123, 2007.

18. Krishna Prasad Pokharel and Ilse Storch. Habitat niche relationships within an assemblage of ungulates in bardia national park, nepal. Acta oecologica, 70:29–36, 2016.

19. Jane Elith and John R Leathwick. Species distribution models: ecological explanation and prediction across space and time. Annual review of ecology, evolution, and systematics, 40, 2009.

20. Richard G Pearson. Species’ distribution modeling for conservation educators and practitioners. Synthesis. American Museum of Natural History, 50:54–89, 2007.

21. Antoine Guisan and Wilfried Thuiller. Predicting species distribution: offering more than simple habitat models. Ecology letters, 8(9):993–1009, 2005.

22. The Global Biodiversity Information Facility (2019) What is GBIF?, howpublished = https://www.gbif.org/what-is-gbif, note = Accessed: 2019-04-03.

23. Jan Beck, Liliana Ballesteros-Mejia, Peter Nagel, and Ian J Kitching. Online solutions and the ‘w allacean shortfall’: what does gbif contribute to our knowledge of species’ ranges? Diversity and Distributions, 19(8):1043–1050, 2013.

24. John Wieczorek, David Bloom, Robert Guralnick, Stan Blum, Markus Döring, Renato Giovanni, Tim Robertson, and David Vieglais. Darwin core: An evolving community-developed biodiversity data standard. PLOS ONE, 7(1):1–8, 01 2012. doi: 10.1371/journal.pone.0029715.

25. GBIF Secretariat. Gbif backbone taxonomy (2019). https://www.gbif.org/species. Accessed: 2018-10-20.

26. Don E Wilson and DeeAnn M Reeder. Mammal species of the world: a taxonomic and geographic reference, volume 1. JHU Press, 2005.

27. Integrated Taxonomic Information System.

28. F Miranda, A Bertassoni, and AM Abba. Myrmecophaga tridactyla. the iucn red list of threatened species 2014: e. t14224a47441961, 2014.

29. Niels Raes and Jesús Aguirre-Gutiérrez. A Modeling Framework to Estimate and Project Species Distributions in Space and Time. 01 2018.

30. R Core Team et al. R: A language and environment for statistical computing. 2013.

31. Alexander Zizka, Daniele Silvestro, Tobias Andermann, Josué Azevedo, Camila Duarte Ritter, Daniel Edler, Harith Farooq, Andrei Herdean, María Ariza, Ruud Scharn, et al. Co-ordinatecleaner: standardized cleaning of occurrence records from biological collection databases. Methods in Ecology and Evolution.

32. Stephen E Fick and Robert J Hijmans. Worldclim 2: new 1-km spatial resolution climate surfaces for global land areas. International journal of climatology, 37(12):4302–4315, 2017.

33. Pascal O Title and Jordan B Bemmels. Envirem: an expanded set of bioclimatic and topographic variables increases flexibility and improves performance of ecological niche modeling. Ecography, 41(2):291–307, 2018.

34. Robert J Hijmans, Jacob van Etten, Joe Cheng, Matteo Mattiuzzi, Michael Sumner, Jonathan A Greenberg, Oscar Perpinan Lamigueiro, Andrew Bevan, Etienne B Racine, Ashton Shortridge, et al. Package ‘raster’. R package, 2015.

35. Wei Shangguan, Yongjiu Dai, Qingyun Duan, Baoyuan Liu, and Hua Yuan. A global soil data set for earth system modeling. Journal of Advances in Modeling Earth Systems, 6(1): 249–263, 2014.

36. Bohdan Slavík and Margaret S Jarvis. Methods of studying plant water relations. 1974.

37. Michael J Crawley. Plant ecology defended. Trends in ecology & evolution, 2(10):304, 1987.

38. Yasmin Hageer, Manuel Esperón-Rodríguez, John B Baumgartner, and Linda J Beaumont. Climate, soil or both? which variables are better predictors of the distributions of australian shrub species? PeerJ, 5:e3446, 2017.

39. Jane Elith, Steven J Phillips, Trevor Hastie, Miroslav Dudík, Yung En Chee, and Colin J Yates. A statistical explanation of maxent for ecologists. Diversity and distributions, 17(1): 43–57, 2011.

40. Steven J Phillips, Robert P Anderson, and Robert E Schapire. Maximum entropy modeling of species geographic distributions. Ecological modelling, 190(3-4):231–259, 2006.

41. Marcelo F Tognelli, Sergio A Roig-Junent, Adriana E Marvaldi, Gustavo E Flores, and Jorge M Lobo. An evaluation of methods for modelling distribution of patagonian insects. Revista chilena de historia natural, 82(3), 2009.

42. James Conolly, Katie Manning, Sue Colledge, Keith Dobney, and Stephen Shennan. Species distribution modelling of ancient cattle from early neolithic sites in sw asia and europe. The Holocene, 22(9):997–1010, 2012.

43. Jane Elith, Catherine H Graham, Robert P Anderson, Miroslav Dudík, Simon Ferrier, Antoine Guisan, Robert J Hijmans, Falk Huettmann, John R Leathwick, Anthony Lehmann, et al. Novel methods improve prediction of species’ distributions from occurrence data. Ecography, 29(2):129–151, 2006.

44. Mary Suzanne Wisz, RJ Hijmans, Jin Li, A Townsend Peterson, CH Graham, Antoine Guisan, and NCEAS Predicting Species Distributions Working Group. Effects of sample size on the performance of species distribution models. Diversity and distributions, 14(5): 763–773, 2008.

45. Carsten F Dormann, Jane Elith, Sven Bacher, Carsten Buchmann, Gudrun Carl, Gabriel Carré, Jaime R García Marquéz, Bernd Gruber, Bruno Lafourcade, Pedro J Leitão, et al. Collinearity: a review of methods to deal with it and a simulation study evaluating their performance. Ecography, 36(1):27–46, 2013.

46. Boris Leroy, Christine N Meynard, Céline Bellard, and Franck Courchamp. virtualspecies, an r package to generate virtual species distributions. Ecography, 39(6):599–607, 2016.

47. RJ Hijmans, S Phillips, J Leathwick, and J Elith. Species distribution modeling with r. r package version 0.8-11, 2013.

48. Yoan Fourcade, Jan O Engler, Dennis Rödder, and Jean Secondi. Mapping species distributions with maxent using a geographically biased sample of presence data: a performance assessment of methods for correcting sampling bias. PloS one, 9(5):e97122, 2014.

49. John A Swets, Robyn M Dawes, and John Monahan. Better decisions through science. Scientific American, 283(4):82–87, 2000.

50. Niels Raes and Hans ter Steege. A null-model for significance testing of presence-only species distribution models. Ecography, 30(5):727–736, 2007.

51. Ben G Holt, Jean-Philippe Lessard, Michael K Borregaard, Susanne A Fritz, Miguel B Araújo, Dimitar Dimitrov, Pierre-Henri Fabre, Catherine H Graham, Gary R Graves, Knud A Jønsson, et al. An update of wallace’s zoogeographic regions of the world. Science, 339 (6115):74–78, 2013.

52. Sylvain Dolédec, Daniel Chessel, and Clémentine Gimaret-Carpentier. Niche separation in community analysis: a new method. Ecology, 81(10):2914–2927, 2000.

53. Andrew J Helmstetter, Tom JM Van Dooren, Alexander ST Papadopulos, Javier Igea, Armand M Leroi, and Vincent Savolainen. Trait evolution and historical biogeography shape assemblages of annual killifish. bioRxiv, page 436808, 2018.

54. Stéphane Dray, Anne-Béatrice Dufour, et al. The ade4 package: implementing the duality diagram for ecologists. Journal of statistical software, 22(4):1–20, 2007.

55. John C Gower. A general coefficient of similarity and some of its properties. Biometrics, pages 857–871, 1971.

56. Martin Maechler, Peter Rousseeuw, Anja Struyf, Mia Hubert, and Kurt Hornik. Package ‘cluster’. Dosegljivo na, 2013.

57. Thomas W Schoener. The anolis lizards of bimini: resource partitioning in a complex fauna. Ecology, 49(4):704–726, 1968.

58. D Rödder and JO Engler. Quantitative metrics of overlaps in grinnellian niches: advances and possible drawbacks. Global Ecology and Biogeography, 20(6):915–927, 2011.

59. Robert Muscarella, Peter J Galante, Mariano Soley-Guardia, Robert A Boria, Jamie M Kass, María Uriarte, and Robert P Anderson. Enm eval: An r package for conducting spatially independent evaluations and estimating optimal model complexity for maxent ecological niche models. Methods in Ecology and Evolution, 5(11):1198–1205, 2014.

60. Emmanuel Paradis, Julien Claude, and Korbinian Strimmer. Ape: analyses of phylogenetics and evolution in r language. Bioinformatics, 20(2):289–290, 2004.

61. Olaf RP Bininda-Emonds, Marcel Cardillo, Kate E Jones, Ross DE MacPhee, Robin MD Beck, Richard Grenyer, Samantha A Price, Rutger A Vos, John L Gittleman, and Andy Purvis. The delayed rise of present-day mammals. Nature, 446(7135):507, 2007.

62. Lam Si Tung Ho, Cecile Ane, Robert Lachlan, Kelsey Tarpinian, Rachel Feldman, Qing Yu, Wouter van der Bijl, and Maintainer Lam Si Tung Ho. Package ‘phylolm’. 2018.

63. Hirotugu Akaike. A new look at the statistical model identification. In Selected Papers of Hirotugu Akaike, pages 215–222. Springer, 1974.

64. Anne E Goodenough, Adam G Hart, and Richard Stafford. Regression with empirical variable selection: description of a new method and application to ecological datasets. PLoS One, 7(3):e34338, 2012.

65. Caroline Grigson. The craniology and relationships of four species of bos: 4. the relationship between bos primigenius boj. and b. taurus l. and its implications for the phylogeny of the domestic breeds. Journal of Archaeological Science, 5(2):123–152, 1978.

66. Laurent Augustin, Carlo Barbante, Piers RF Barnes, J Marc Barnola, Matthias Bigler, Emiliano Castellano, Olivier Cattani, Jerome Chappellaz, Dorthe Dahl-Jensen, Barbara Del-monte, et al. Eight glacial cycles from an antarctic ice core. Nature, 429:623–628, 2004.

